# Prime editing enables high-throughput identification and characterization of selective MutSβ inhibitors

**DOI:** 10.64898/2026.10.05.755959

**Authors:** Yangwode Jing, Jun Yan, Denny Yang, Yanjin Chen, Issam Senoussi, James Woodward, Katya Williams, Hahn Kim, Kangning He, Herwin Daub, Vinay Dahiya, Maren Thomsen, Jia Ling, Purnima Ravisankar, Yuri Pritykin, Gabriel Balmus, Petr Cejka, Jeffrey A. Hussmann, Longbin Liu, Thomas F. Vogt, Michael Finley, Britt Adamson

## Abstract

DNA mismatch repair (MMR) maintains genome stability but paradoxically drives expansion of pathogenic repeats responsible for trinucleotide repeat disorders. Attenuating repeat instability through targeted inhibition of MMR is therefore a therapeutic strategy of interest; however, progress is constrained by a lack of robust methods for defining and isolating MMR subpathways. We exploited features of prime editing to profile the effects of core MMR genes on thousands of programmed DNA mispairs in cells and resolved distinct MMR processes, including a noncanonical “loop-directed” mechanism. Guided by these results, we built a set of scalable reporters that detect loss of separable MMR processes and screened ∼72,000 small molecules for activity against MutSβ-directed MMR. This effort identified a novel series of MutSβ inhibitors that reduced (CAG)_n_ expansion in a cell model. Altogether, this study establishes several methods for systematically interrogating MMR, with demonstrated capacity to identify inhibitors of therapeutic value.

## INTRODUCTION

Huntington’s disease (HD) is an incurable, neurodegenerative disorder characterized by progressive motor dysfunction, cognitive decline, and psychiatric disturbances, leading to severe disability and death.^1^ HD exhibits a pattern of autosomal dominant inheritance linked to an expanded trinucleotide CAG repeat in exon 1 of the gene *huntingtin* (*HTT*),^2^ placing it within a broader class of disorders caused by unstable repeats in the human genome.^3^ As defined in a recent position paper,^4^ individuals with ≥40 CAG repeats in *HTT* have highly penetrant HD (>98%), with >60% probability of clinical motor diagnosis by age 65 (at a diagnostic confidence level of 99%), whereas individuals born with more moderately expanded *HTT* alleles (36-39 CAGs) show reduced penetrance and delayed progression of the disease. Of significance, disease-causing CAG tracts undergo additional lengthening in select cell types of people with HD, and somatic expansion of these alleles in striatal medium spiny neurons is thought to contribute to neurodegeneration and disease progression.^5,6^ Halting or slowing somatic lengthening of repeats has thus emerged as a therapeutic strategy for HD and other repeat expansion disorders of unmet medical need.

Two complementary lines of evidence have established a tractable path forward for developing trinucleotide repeat-stabilizing interventions: (1) genome-wide association studies have identified genetic modifiers of somatic (CAG)_n_ expansion and HD clinical phenotypes in and around genes associated with DNA mismatch repair (MMR),^7–13^ with some variants able to delay disease onset or progression, and (2) disruption of several of those modifier genes has been shown to attenuate (CAG)_n_ expansion in experimental models of HD, including cultured cells,^14,15^ patient-derived stem cells and differentiated neurons,^16,17^ and mice.^18–22^ Together, these findings implicate mechanisms of MMR in driving somatic instability of HD-causing alleles and suggest that targeted inhibition of select MMR components could provide therapeutic benefit.

Although implicated in somatic repeat instability, the primary role of MMR is to maintain genome stability by correcting base-base mismatches and small loop-out structures that arise during normal cellular processes (e.g., DNA replication and transcription).^23,24^ Exactly how MMR goes awry to expand unstable CAG repeats remains unclear; however, converging lines of evidence implicate MutSβ-directed mechanisms. For context, small DNA mispairs can be detected by one of two heterodimeric MMR sensor complexes, MutSα (MSH2-MSH6) or MutSβ (MSH2-MSH3). These sensors bind overlapping but distinct classes of DNA mispairs and recruit downstream factors to identify and remove error-containing DNA strands (Figure 1A). Supporting a role for MutSβ in somatic repeat expansion, the gene encoding the unique subunit of MutSβ (*MSH3*) is a consistent modifier of HD across human genetic studies and promotes (CAG)_n_ expansion in HD models. Moreover, MutSβ binds with high affinity to (CAG)_n_ loop-out structures,^25–27^ which can arise within tracts of CAG repeats due to strand slippage and may initiate repeat expansion processes. MutSα, by contrast, has been reported to have a minimal^19^ or even opposite role^18^ in somatic repeat instability.

**Figure 1.**
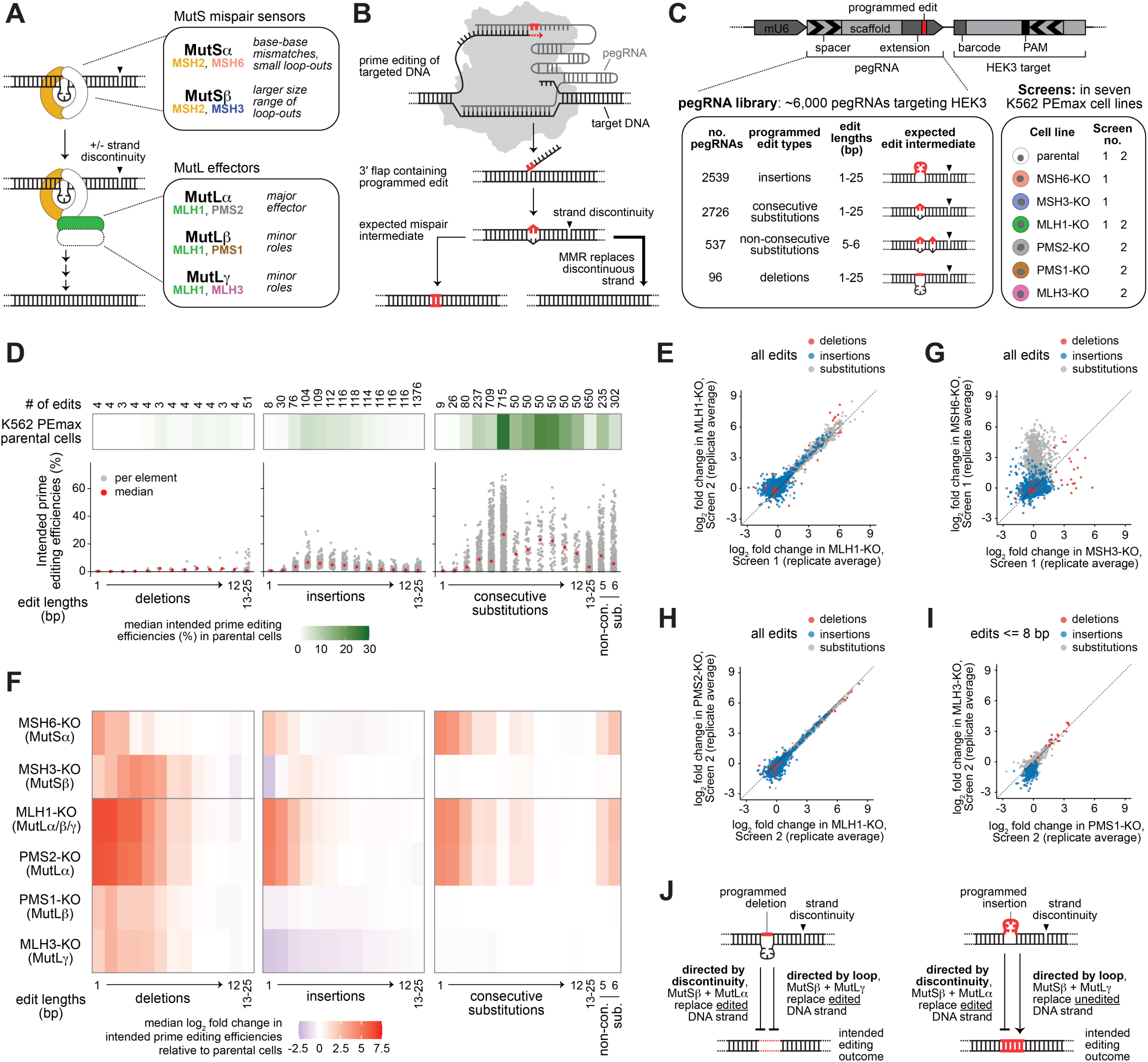
Pooled prime editing screens map mismatch repair gene functions across different prime edit types. (A) Schematic of DNA mismatch repair. Mismatches are sensed by MutSα/β sensor complexes, which recruit MutLα/β/γ and other effectors to complete downstream repair processes. Black triangles indicate a strand discontinuity in 1A-1C. (B) Simple model of prime editing and programmed mispair rejection by DNA mismatch repair. (C) Schematic depicting prime editing library using the HEK3 target site and related screens. Self-targeting lentiviral constructs (top) were used for delivery of the target sequence and pegRNAs programming many DNA mispairs to that site (bottom left). Screens were performed in K562 PEmax parental cells and derivative cell lines with core MMR genes knocked out (bottom right). Arrows within the depicted construct indicate directionality of the pegRNA spacer and matched protospacer within the target. Details on the other library and screen are included in Figure S1B. (D) Heatmaps (top) show median intended prime editing efficiencies at the HEK3 target site in K562 PEmax parental cells grouped based on edit length (1-25 bp, with 13-25 bp edits collapsed into one group) and type (deletions, insertions, consecutive substitutions, and non-consecutive substitutions, which are abbreviated as non-con. sub.). The number of prime edits in each group is included. Dot plots (bottom) show individual (grey dots) and median (red dots) intended prime editing efficiencies within each group. (E) Comparison of log_2_ fold changes in intended prime editing efficiencies at the HEK3 target site in MLH1-KO cells between two different screens (HEK3 Screens 1 and 2). Each dot represents an element in the prime editing library, colored by edit type (red, deletions; blue, insertions; grey, substitutions). Black dashed line indicates y = x. (F) Heatmaps show median log_2_ fold changes in intended prime editing efficiencies for each knockout cell line at the HEK3 target site, stratified by prime edit length and type, as in D. Grey horizontal lines between heatmaps separate results from HEK3 Screens 1 and 2. (G) Comparison of log_2_ fold changes in intended prime editing efficiencies at the HEK3 target site in MSH3-KO cells (x-axis) to those from MSH6-KO cells (y-axis). Each dot represents an element in the prime editing library, colored by edit type, as in E. Black dashed line indicates y = x. (H) Comparison of log_2_ fold changes in intended prime editing efficiencies at the HEK3 target site in MLH1-KO cells (x-axis) to those from PMS2-KO cells (y-axis). Each dot represents an element in the prime editing library, colored by edit type, as in E. MLH1-KO data was from HEK3 Screen 2. Black dashed line indicates y = x. (I) Comparison of log_2_ fold changes in intended prime editing efficiencies at the HEK3 target site in PMS1-KO cells (x-axis) to those from MLH3-KO cells (y-axis). Each dot represents an element in the prime editing library, colored by edit type, as in E. Only edits <= 8 bp were included in this analysis. Black dashed line indicates y = x. (J) Schematics of MMR pathways that can act on loop-out intermediates arising during deletion or insertion prime editing: discontinuity-directed MMR mediated by MutSβ+MutLα, and loop-directed MMR mediated by MutSβ+MutLγ. The former suppresses installation of both deletion and insertion prime edits, while the latter suppresses deletions and promotes insertions.

MutSβ thus represents a compelling drug target; however, given the central role of MMR in genome maintenance, concomitant risk of elevated cancer susceptibility is a concern. Reinforcing this concern, inherited mutations in several MMR genes (e.g., *MSH6* and *MSH2*) predispose individuals to cancer^28,29^ and mutations in *MSH3* have shown associations with cancer and adenomatous polyposis.^30–32^ Evidence supporting a causal role for MutSβ dysfunction in development of cancer though is lacking, and multiple studies have shown that loss of *MSH3* does not elevate tumor incidence or affect survival in mice, unlike disruption of other MMR genes.^33,34^ Additionally, knocking down *MSH3* with antisense oligonucleotides or siRNAs does not alter DNA damage response, cancer-related pathways, or tumor-associated microsatellite instability.^16,21^ MutSβ inhibition may therefore have a tolerable safety profile, and current evidence supports investigating that possibility, whilst also underscoring a need for pathway-selective inhibition strategies.

Despite the therapeutic potential of MutSβ inhibition, two major barriers exist: (1) we lack small molecules with selective activity against MutSβ confirmed in cells, and (2) we lack robust cellular assays for identifying and characterizing such compounds. The latter challenge represents a particularly significant bottleneck and motivates the development of new methods for measuring MutSβ deficiency in cells, to accelerate inhibitor discovery and improve characterization of the few inhibitors already identified.^35,36^ Notably, a common limitation of existing methods for assaying MutSβ function is reliance on repeat expansions as endpoint readouts.^37–41^ This design limits the dynamic range of detection relative to background, imposes a need for long periods of cell growth before data collection, and can make the assays difficult to implement. Moreover, assays based on repeat expansion do not always fully resolve the individual contributions of MutSα and MutSβ to observed phenotypes. By contrast, *in vitro* biochemical assays provide a way to rapidly measure MutSβ activity, often with well-resolved signals;^42–45^ however, these assays lack cellular context and thus do not account for the influence of other cellular pathways or capture indirect effects.

Here, we present a suite of cell-based assays that enable reliable, rapid, and selective detection of MutSβ inhibition. To develop these assays, we profiled prime editing efficiencies across thousands of edits in cells with targeted disruption of core MMR genes. These screens distinguished distinct MMR activities, including those coordinated by the MutSα and MutSβ sensor complexes, and resolved a poorly characterized, noncanonical mechanism of “loop-directed” MMR. Prime edit types found to be selectively suppressed by MutSα or MutSβ were then repurposed to build a set of highly scalable, sensor-selective MMR reporters. We used one of these reporters to screen ∼72,000 small molecules for activity against MutSβ and identified a novel series of MutSβ inhibitors. Characterization of these inhibitors revealed selectively for MutSβ, with no observed activity against MutSα in cells, and demonstrated reduction of (CAG)_n_ expansion in a cellular model of somatic instability. Together, our results establish a set of versatile tools for studying MMR and for advancing small molecule inhibitors of MutSβ towards clinical applications in repeat expansion diseases.

## RESULTS

### Systematic evaluation of MutS and MutL activities in MMR using prime editing

Prime editing is a CRISPR-based genome editing technology that can write small insertions, deletions, and substitutions into genomes using reverse transcription.^46^ We and others have shown that prime editing is suppressed by MMR, with inhibition occurring in accordance with known MMR substrate criteria.^47–50^ That is, MMR suppresses prime editing when programmed edits are expected to form MMR substrate-mimicking intermediates (e.g., short runs of base mispairs, extrusions of 1-10 nt^51–55^) but has generally less impact when intermediary editing structures are expected to be poor MMR substrates (e.g., C-C mispairs introduced by G to C edits^51,56,57^ or longer extrusions of >10 nt^53,54^). Collectively, these observations have established a model of prime editing wherein transient DNA mispairs are generated during edit installation and recognition of those mispairs by MMR leads to edit removal (Figure 1B).

The substrate recognition criteria of the two canonical MutS sensors in humans (MutSα and MutSβ) have been well described. MutSα detects mispaired DNA bases and small DNA loop-outs (∼1-2 nt),^58,59^ whereas MutSβ recognizes larger loops (∼1-15 nt) but not base-base mispairs (Figure 1A).^42,59,60^ However, how these preferences, which have been defined largely by biochemical analyses, stratify across mispairs in the cellular environment has been minimally described.^61^ Moreover, how three MutL effector complexes (MutLα/β/γ) coordinate repair downstream of the sensors is incompletely understood, particularly with respect to repeat instability in neurons.^62–64^ We reasoned that, by profiling the effects of MutS and MutL genes across many prime edits, each designed to generate a different DNA mispair, we could elucidate separable MMR subpathways (i.e., those directed by different classes of MMR substrates) (Figure S1A).

To test this idea, we generated two prime editing libraries, each encoding a single target sequence (HEK3 or *FANCF*) and 6000 prime editing guide RNAs (5898 and 5922 edit-specifying pegRNAs and 102 and 78 non-editing control pegRNAs, respectively) and each designed to generate thousands of different edits, and thus thousands of distinct DNA mispair structures, in the corresponding target site (Figures 1C and S1B; Table S1). Sequencing these libraries detected >99% of the pegRNAs, 90% of which were within a range of < 4-fold (Figure S1C). After characterizing these libraries, we transduced them into K562 cells expressing a standard prime editor (PEmax)^51,65^ and several derivative knockout (KO) cell lines, each with genetic disruption of one of six MMR genes: *MSH6*, *MSH3*, *MLH1*^56^, *PMS2*, *PMS1*, or *MLH3* (Figures S1D-S1N; Tables S2-S4). These cell lines were transduced at low multiplicities of infection (0.25-0.45) in one of two screening groups: (1) parental, MSH6-KO, MSH3-KO, and MLH1-KO cell lines for one screen with our HEK3-targeted library (HEK Screen 1) and one with our *FANCF*-targeted library (FANCF Screen), or (2) parental, MLH1-KO, PMS2-KO, PMS1-KO, and MLH3-KO cell lines for a second HEK3 screen

(HEK3 Screen 2) (Figures 1C and S1B). Cells were selected for library integration, and after seven days of growth, we quantified intended editing efficiencies by sequencing pegRNA-target pairs and calculating the percentage of corresponding target reads harboring the programmed prime edit (Tables S5 and S6).

Prior to any other analyses, we evaluated the reproducibility and validity of the intended editing efficiencies measured in our screens. Indicating low technical variability and high reproducibility, replicate intended editing efficiencies were well correlated (Figure S2A), even when measured from non-concurrent screens (i.e., conditions repeated in HEK3 Screens 1 and 2). Moreover, when stratified by edit length and edit type, intended editing efficiencies from MMR-proficient parental cells confirmed two observations from previous studies (Figures 1D and S2B): (1) substitutions of ≥3 contiguous base pairs (bps) were installed more efficiently than single substitutions or those spanning 2 bps,^51^ even when restricting comparisons to edits that did not disrupt the PAM sequence (8.1-fold increase for 3-4 bp consecutive substitutions in HEK3, *p* < 0.0001, and 2.1-fold increase for those substitutions in *FANCF*, *p* < 0.0001), and (2) small insertions were installed more efficiently than deletions of similar size.^66^ The first of these observations has been attributed to poor MutS recognition of contiguous mismatches,^51^ while the second has yet to be explained.

We next quantified the impact of *MSH6*, *MSH3*, *MLH1*, *PMS2*, *PMS1*, and *MLH3* on edits evaluated in our screens by calculating log_2_ fold changes in intended editing efficiencies between K562 PEmax parental cells and the knockout cell lines. These measurements were also reproducible between replicates (Figures 1E and S2C) and captured expected features (Figures 1F and S2D-S2F). For example, disruption of *MLH1*, which causes loss of all three MutL complexes and thus broadly impedes MMR (Figures S1M and S1N), increased all types of small prime edits (∼1-8 bps) but had generally less effect on longer edits (>8 bps).^51,53^ Additionally, loss of each of the two MutS sensors in MSH6-KO and MSH3-KO cells impacted distinct groups of edits (Figure 1G); that is, we observed an increase in small insertions, small deletions, and substitutions (consecutive and non-consecutive) in MSH6-KO cells, which lack expression of MutSα but not MutSβ, and an increase in deletions across a wider range of lengths, some insertions, and generally not substitutions in MSH3-KO cells, which lack expression of MutSβ but not MutSα (Figures 1F, S1H, S1I, and S2D-S2F). These results agree with prior examination of a limited number of prime edits in *MSH6* and *MSH3* knockout cells^54^ and, alongside expected mispair intermediates for each stratified edit type (Figures 1C and S1B), reinforce known MutS substrate recognition rules (Figure 1A).

We next considered results from our MutL knockout cell lines: MLH1-KO, PMS2-KO, PMS1-KO, and MLH3-KO. Although eukaryotic cells express multiple MutL homologs, MutLα is considered the major effector of canonical MMR,^67^ which occurs in five steps: (1) mispair recognition by MutSα or MutSβ, (2) recruitment of the MutLα complex, (3) MutLα-catalyzed nicking of the error-containing DNA strand, which is discriminated from the “correct” DNA strand by a preexisting strand discontinuity or nick somewhere adjacent to the mispair, (4) excision of the error-containing DNA, and (5) DNA resynthesis.^23,68^ In our screens, PMS2-KO cells, which lack MutLα but not the other MutL complexes, demonstrated the strongest and most consistent promotion of prime editing among MutL-specific knockout cell lines, and generally phenocopied MLH1-KO cells, which lack all three MutL complexes (Figure 1H). These results confirm a major role for MutLα-catalyzed MMR in rejecting prime edited DNA, likely through excision of the inherently discontinuous, reverse-transcribed flaps or from interference with flap annealing^51^ (Figure 1B).

Of significance, our screen data also revealed several unanticipated results. For example, we found that the editing efficiencies of 1 bp insertions were decreased in MSH3-KO cells, relative to parental K562 PEmax cells, and editing efficiencies of 2 bp insertions were minimally altered in those cells (Figures 1F and S2D-S2F). According to the current model of prime editing (Figure 1B), which evokes only canonical MMR, those edits should be more efficiently installed without MutSβ, similar to effects observed for slightly longer insertions (e.g., 3-6 bp) and deletions of the same size (1-2 bp). Additionally, we found that PMS1-KO and MLH3-KO cells, which lack MutLβ and MutLγ, respectively, produced intriguing phenotypes; that is, we observed increased editing efficiencies for small deletions and, surprisingly, decreased editing efficiencies for similarly sized insertions (∼1-8 bp) (Figure 1I).

Together, these unexpected results suggest that a noncanonical mechanism of “loop-directed” MMR, which was recently described *in vitro*,^69,70^ acts on some prime edit types. In this mechanism, MutSβ-stimulated MutLγ cleaves DNA across from small loop-out structures to retain extruded DNA in repaired duplexes. This mechanism should, accordingly, suppress small deletion edits but have opposite effects on small insertions (Figure 1J). Isolated loss of this mechanism in MLH3-KO cells, which lack the relevant endonuclease, MutLγ, provides a compelling explanation for the deletion and insertion phenotypes observed in those cells (Figures 1I and S2D). Moreover, parallel activity of this mechanism alongside canonical MMR explains the idiosyncratic insertion phenotypes observed in cells lacking MutSβ (i.e., MSH3-KO cells). To elaborate, if loop-directed MMR acts on loop-out structures formed during editing, those intermediates should be processed by one of two MutSβ-driven MMR pathways: (1) MutLα-catalyzed removal of the reverse transcribed DNA (i.e., canonical MMR) or (2) MutLγ-catalyzed removal of the strand opposite the loop. These pathways should therefore both suppress small deletion edits (Figure 1J, left), but we expect them to exert opposing effects on small insertions, due to different positioning of the loop-out relative to the reverse transcribed flap (Figure 1J, right). Under this dual-pathway model, loss of MutSβ would consistently increase small deletion editing, as observed, but insertion phenotypes would likely vary, depending on the relative activity of each subpathway at the initiating substrate. Our data thus suggest an expanded model of prime editing, in which installation of small insertions, but not deletions, is aided by an endogenous repair mechanism. This model explains why small insertion edits are more efficiently installed than small deletions in MMR-proficient cells^66^ (Figures 1D and S2B). Additionally, these results show that, despite complex interactions among MMR genes, prime editing can resolve distinct MMR pathways.

### Development of MutSα/β-specific reporters using different prime edit types

Our genetic screens showed that prime edits can differentially engage distinct MMR activities, with several edit types able to distinguish MutSα- and MutSβ-driven MMR. For example, most 1-5 bp substitutions were suppressed by *MSH6* but not *MSH3*, while 4-6 bp deletions had reciprocal specificities (Figures 1F and S2D-S2F). Further evaluation of 1 bp substitutions and 5 bp deletions across additional target site sequences in K562 cells confirmed that loss of *MSH6* but not *MSH3* consistently increased installation of the former edit type (∼2-14 fold) whereas *MSH3* but not *MSH6* had an inhibitory effect on the later (∼2-10 fold) (Figure 2A). Similar results from two additional cell lines (i.e., HAP1 and RPE-1 hTERT cells) suggest that these sensitivities reflect intrinsic, sensor-specific stimulation of MMR, rather than cell line-specific effects (Figure S3). Moreover, confirming that these phenotypes are not artifacts of clonal selection, ectopic expression of MSH6 in K562 and HAP1 MSH6-KO cells reduced 1 bp substitution editing, effectively restoring suppression towards levels observed in parental cells, and ectopic expression of MSH3 in K562 and HAP1 MSH3-KO cells showed similar effects on a 5 bp deletion (Figures 2B and 2C, Methods). These results establish that 1 bp substitutions and 5 bp deletions can robustly, specifically, and consistently stimulate the activity of only one MMR sensor complex, making their editing efficiencies reliable readouts of sensor-directed MMR.

**Figure 2.**
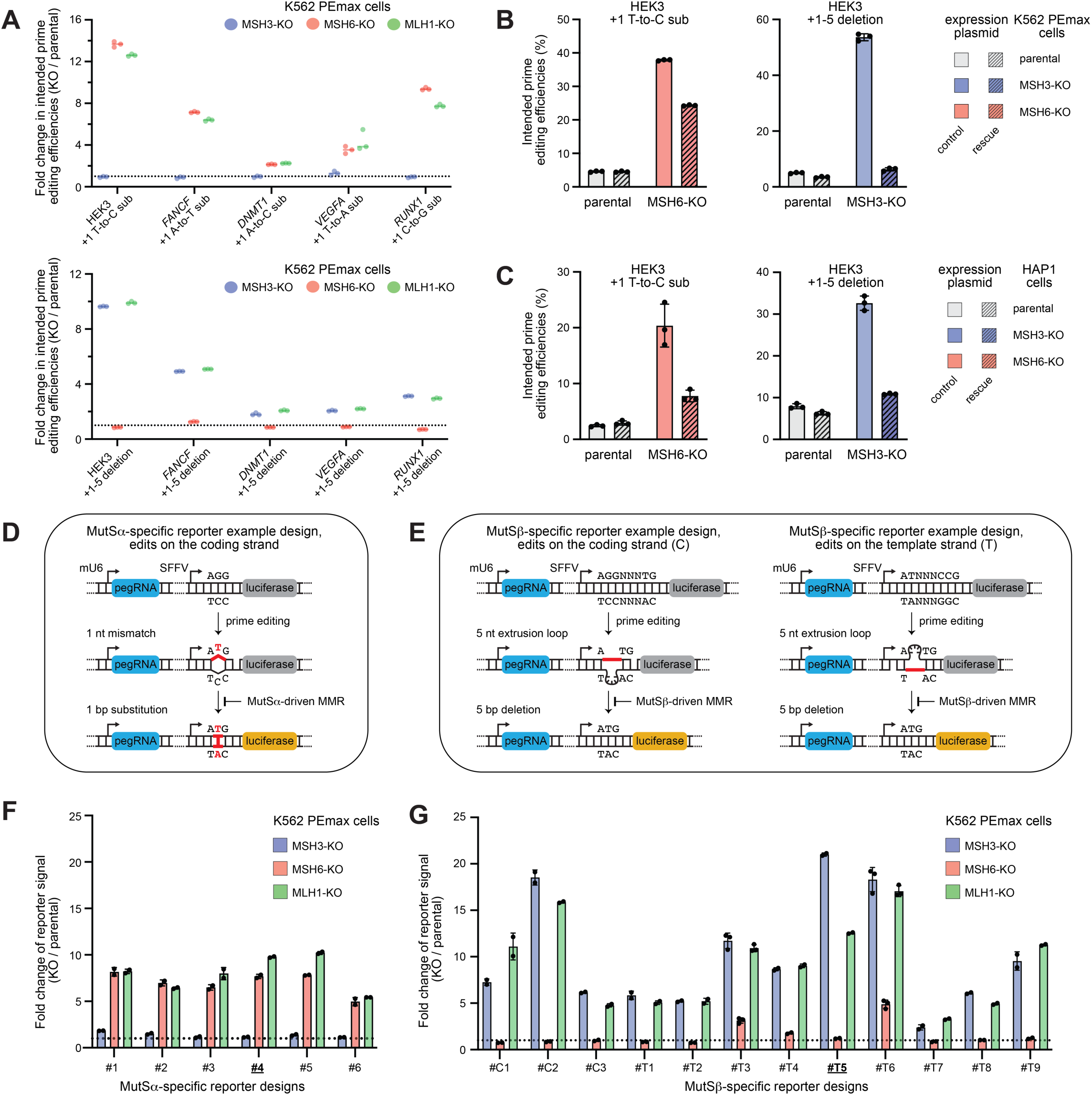
MutSα/β-specific MMR reporters engineered with MutS resolving prime edits. (A) Fold changes in intended prime editing efficiencies for 1 bp substitution edits (top) or 5 bp deletion edits (bottom) at indicated genomic loci in indicated cell lines relative to parental cells. Groups of dots represent data from n = 3 biological replicates. pegRNA plasmids were delivered by nucleofection. (B) Intended prime editing efficiencies at the endogenous HEK3 target site in K562 PEmax parental cells or indicated knockout cell lines, after nucleofection of pegRNA and rescue or control expression plasmids. Bars represent means of n = 3 biological replicates and error bars indicate standard deviations. (C) Intended prime editing efficiencies at the endogenous HEK3 target site in HAP1 parental cells or indicated knockout cell lines, after nucleofection of pegRNA, PEmax editor, and rescue or control expression plasmids. Bars represent means of n = 3 biological replicates and error bars indicate standard deviations. (D) Schematic of MutSα-specific reporters. (E) Schematics of MutSβ-specific reporters. Edits were designed to target either the coding strand (C, left) or the template strand (T, right) of the luciferase indicator gene. (F-G) Characterization of MutSα-specific reporters (F) and MutSβ-specific reporters (G) in K562 PEmax cells. Reporters were delivered to K562 PEmax parental cells or indicated knockout cell lines by lentiviral transduction. Dots represent fold changes in luminescence, bars represent means of n = 2 or 3 technical replicates, and error bars indicate standard deviations. Reporters selected for further characterization are indicated **<u>in bold and underlined</u>** (#4 for MutSα- and #T5 for MutSβ-specific reporters).

We next leveraged these sensor-selective edit classes to build a series of reporters that detect disruption of either MutSα- or MutSβ-directed MMR. We envisioned reporters in which installation of a MutSα- or MutSβ-suppressed edit would install an in-frame start codon and thus turn on a detectable gene product (Figures 2D and 2E). Elevated editing upon inhibition or loss of the corresponding MMR sensor would, in turn, increase expression of the indicator gene (herein, luciferase) and signal loss of that sensor. Unlike many alternative MMR assays,^71–76^ these reporters would be delivered with lentiviral transduction and would thus monitor MMR occurring directly in the genome. Notably, to detect delivery, our reporters would also constitutively express a marker of transduction from the same mRNA as the indicator gene, driven by an internal ribosome entry site (IRES),^77^ similar to a system we used previously^65^ (Methods).

To build these reporters, we chose eighteen target sequences previously shown to support prime editing^56,78^ and designed eighteen 1 bp substitution and 5 bp deletion edits to introduce an ATG codon or its reverse complement (CAT) into each of those sequences. We cloned pegRNA sequences specifying these edits and the corresponding targets into the same expression constructs, with the targets positioned between a promoter driving mRNA expression and the indicator gene such that the specified edits would generate an in-frame ATG codon on the coding strand (Figures 2D and 2E; Table S7; Methods). We then evaluated each of the resulting “self-targeting” reporters in cells expressing PEmax, measuring induction of luciferase in parental cells and MSH3-KO, MSH6-KO, and MLH1-KO derivatives three days post reporter transduction (Figures 2F and 2G; Table S7). All six constructs designed with 1 bp substitutions were robustly activated in *MSH6* knockout cells but not *MSH3* knockouts. We refer to these as our MutSα-specific reporters. Conversely, all twelve constructs designed with 5 bp deletions showed reciprocal specificities. We refer to these as our MutSβ-specific reporters. Notably, all 18 reporters were activated in cells lacking MLH1, consistent with loss of both MutSα- or MutSβ-driven MMR. Collectively, these findings establish a prime editing-based platform for selectively detecting inhibition of MutSα- and MutSβ-directed MMR.

### Establishing MutSα/β-specific reporters as robust and scalable readouts of sensor-resolved MMR

We selected two reporters for further characterization: MutSα-specific #4 and MutSβ-specific #T5 (Figures 2F and 2G; hereafter referred to without the #4 and #T5 identifiers). Testing these reporters in parental K562 PEmax cells and seven derivative knockout cell lines recapitulated phenotypes expected from our genetic screens; that is, disruption of *MSH3*, *MSH2, MLH1*, *PMS2*, *PMS1*, and *MLH3* activated the MutSβ-specific reporter, but disruption of *MSH6* did not; whereas, disruption of *MSH6*, *MSH2*, *MLH1*, and *PMS2* robustly activated the MutSα-specific reporter, but disruption of *MSH3* and *MLH3* did not (Figure 3A). These phenotypes were stable over consecutive days (Figure S4A), were highly correlated with changes in intended prime editing determined by sequencing (Figure S4B), and were reversible by re-expression of the disrupted genes, with the strength of rescue often scaling with the level of ectopic protein expression (Figures 3B, 3C, and S4C-S4H; Methods). We also found that reporter phenotypes were robust across experimental conditions, as strong effects reproduced with an alternative delivery vector (Figure S4I), an additional cell line (RPE-1 hTERT; Figure S4J), and a distinct method of perturbing MutSβ (siRNA; Figure S4K). Moreover, in addition to pathway disruption, our MutSβ-specific reporter was able to register increased MutSβ-driven MMR, as evident in the reduction of reporter signal observed upon strong overexpression of MSH3 (Figures 3B, S4D, and S4L). These results establish our MutSα/β-specific reporters as robust and reliable indicators of sensor-resolved MMR, amenable to functional studies.

**Figure 3.**
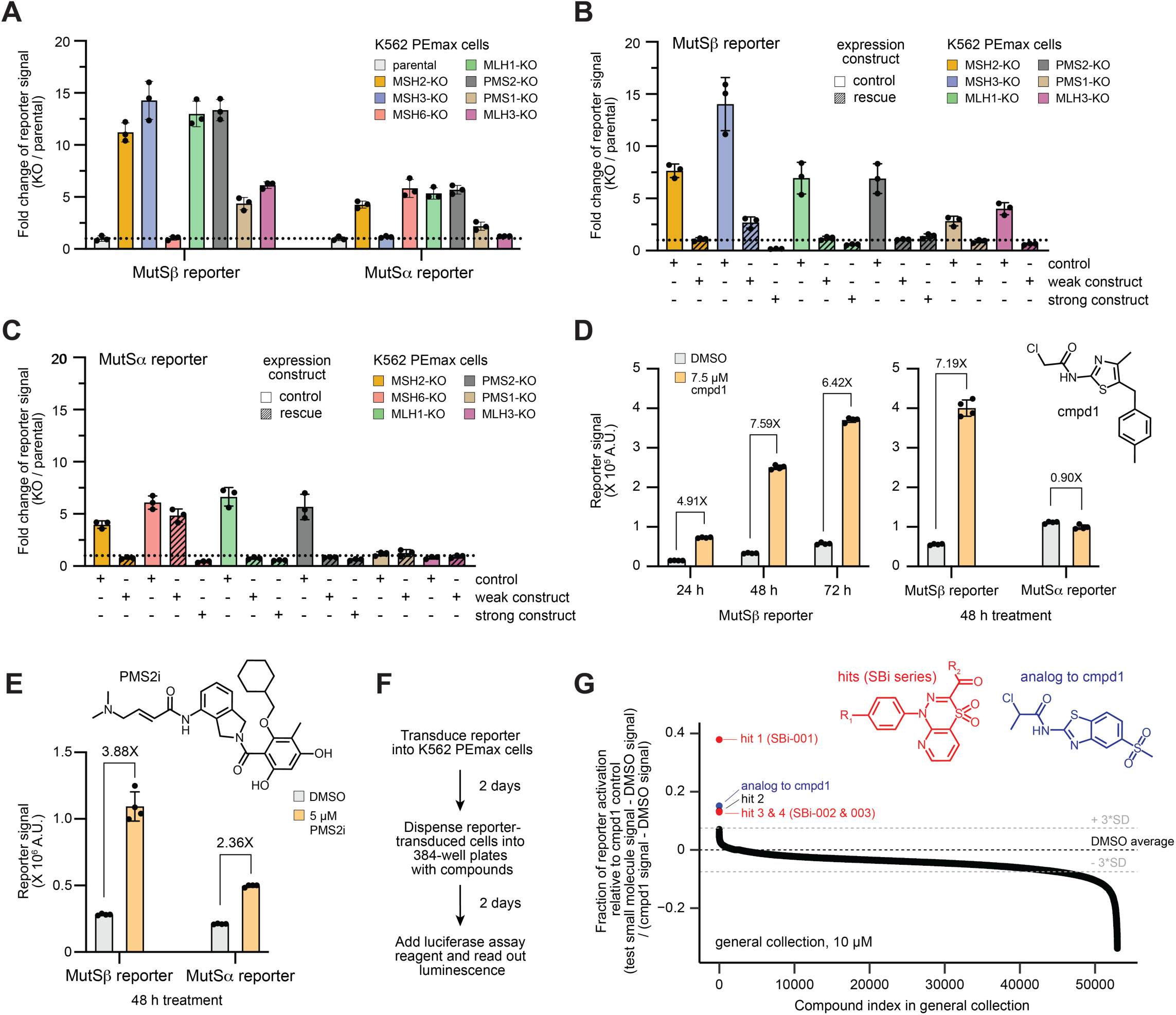
MutSα/β-specific reporters enable sensor-resolved measurement of MMR disruption and high-throughput small molecule screening. (A) Characterization of our primary MutSβ- and MutSα-specific reporters in K562 PEmax parental cells and indicated knockout cell lines. Dots represent fold changes in luminescence, bars represent means of n = 3 biological replicates, and error bars indicate standard deviations. (B) MMR rescue experiments with our primary MutSβ-specific reporter. A weak or strong rescue construct or control (Methods) was delivered to the indicated K562 PEmax knockout cell line by lentivirus, with the rescue constructs matched to the knocked out gene. Dots represent fold changes in luminescence, bars represent means of n = 3 biological replicates, and error bars indicate standard deviations. (C) As in B but for our primary MutSα-specific reporter. (D) MutSβ-specific reporter activation by cmpd1 in reporter-transduced K562 PEmax cells after drug treatments of varying durations (left). Comparison of the effect of cmpd1 on our primary MutSβ- and MutSα-specific reporters in reporter-transduced K562 PEmax cells after 48 hours drug treatments (right). Dots represent raw luminescence signals, bars represent means of n = 4 biological replicates, and error bars indicate standard deviations. cmpd1 structure is depicted (inset). (E) Comparison of the effect of PMS2i on our primary MutSβ- and MutSα-specific reporters in reporter-transduced K562 PEmax cells after 48 hours drug treatments. Dots represent raw luminescence signals, bars represent means of n = 4 biological replicates, and error bars indicate standard deviations. PMS2i structure is depicted (inset). (F) Description of our optimized MutSβ-specific reporter assay used for high-throughput small molecule screening. Luminescence was read out using Nano-Glo luciferase assay. (G) Results from screening the general collection of small molecules using our primary MutSβ-specific reporter (n = 1). Dots represent fractions of reporter activation relative to cpmd1 controls, defined as background-subtracted luminescence from individual wells normalized to the mean of background-subtracted luminescence from on-plate cmpd1 controls, such that 0 represents average results from DMSO controls (black dashed line) and 1 represents average results from cmpd1 controls (Methods). Grey dashed lines indicate 3 standard deviations from the DMSO average (+/- 3*SD). Structures relevant to 4 out of 5 hit molecules (blue and red points, Table S8) are depicted (insets), with R_1_ and R_2_ representing different functional groups for SBi molecules (Figure 4A). We note that data for a few hits overlap on this plot, leaving only 3 such points visible.

As proof-of-concept for chemical applications, we next used our reporters to evaluate two recently described MMR inhibitors: cmpd1,^35^ which has been shown to covalently bind the DNA-binding domain of MSH3 *in vitro* but has unresolved cellular activity,^36^ and a recently disclosed PMS2 inhibitor^79^ (NP1867, hereafter called PMS2i). Indicative of selective MutSβ inhibition, a single dose of cmpd1 (7.5 μM) activated our MutSβ-specific reporter in K562 PEmax cells (with increased signal relative to controls observed as early as 24 hours post treatment), but, under similar conditions, did not activate our MutSα-specific reporter (Figure 3D). Further testing revealed that the effect of cmpd1 on our MutSβ-specific reporter was dose-dependent, with maximal activation observed at concentrations near the IC_50_ for cytotoxicity of cmpd1, beyond which activation of the reporter was diminished, likely due to reduced cell viability (Figures S5A and S5B). These results show that cmpd1 selectively inhibits MutSβ-driven MMR in cultured cells. Moreover, we observed similar activity for cmpd1 in three additional cell lines (HAP1, HeLa, U-2 OS; Figure S5C), supporting a general effect, but as expected, cmpd1 did not activate our reporter in MMR-deficient cells (Lenti-X 293T, HCT 116; Figure S5D). PMS2i, by contrast, activated both of our primary reporters, consistent with MMR inhibition downstream of both MutSα and MutSβ (Figures 3E and S5E).

Having established that our reporters robustly detect chemical inhibition of MMR, we next sought to apply our MutSβ-specific reporter to screen for additional MutSβ inhibitors. Towards that end, we developed a scalable, four-day reporter assay (Figure 3F) wherein K562 PEmax cells transduced with the MutSβ-specific reporter are grown for two days, dispensed into microwell plates containing test and control compounds, and analyzed for luminescence with Nano-Glo luciferase assay reagents two days later. We rigorously optimized the performance of this assay by (1) determining transduction conditions that provide maximal activation (Figure S5F), (2) confirming that NanoLuc provided the largest dynamic range among commonly used luciferases, and establishing conditions for use in both 96- and 384-well plates (Figure S5G). Together, these efforts produced a scalable MutSβ-specific reporter assay for high-throughput, array-based small molecule screening, designed to detect cell-active inhibitors of MutSβ-driven MMR.

### Application of a MutSβ-specific reporter to high-throughput small molecule screening

Using our optimized MutSβ-specific reporter assay (Figure 3F), we screened 71,623 small molecules for MutSβ inhibitors. We screened two collections: one containing 5,923 bioactive molecules with annotated targets and known effects, and another comprising 65,700 molecules with drug-like structures but no annotated targets or effects (organized as 47,780 individual compounds and 4,480 mixtures of stereoisomers). We evaluated the smaller, bioactive collection on 20 microwell plates at two concentrations (10 μM and 20 μM) and the much larger, general collection on 164 plates at only a single concentration (10 μM). On each plate, we included 32 wells of cmpd1 (10 μM) and 32 wells of solvent (DMSO) to serve as positive and negative controls, respectively. Using data from these controls, we calculated per-plate Z′-factors^80^ and, to ensure high data quality, rescreened plates with low Z′-factors (< 0.2, 9 total plates). Indicative of good dynamic range and low variability, our final data set had a mean Z′-factor of 0.558 ± 0.095 (Figure S5H). To quantify inhibition of MutSβ-driven MMR for each compound, we next calculated background-subtracted luminescence for each well and normalized those measurements to on-plate cmpd1 controls, generating relative inhibition scores (Figures 3G and S5I; Methods). Hit molecules were determined as those for which this metric exceeded the DMSO control average by ≥ 3 standard deviations.

From the bioactive collection, we identified 55 hits. Providing confidence in these findings, results from this collection were reproducible across the two concentrations screened (Pearson’s correlation r = 0.86; Figure S5J) and roughly a third of the hits (18) scored at both 10 μM and 20 μM (Table S8). Moreover, one hit from the bioactive collection, the FDA-approved antiviral Ribavirin, scored repeatedly from multiple collection entries (i.e., three times at 10 μM and twice at 20 μM). Additionally, retesting the strongest hits from this collection (i.e., 28 with > 0.1 normalized reporter activation) confirmed results for 11 molecules, albeit often with a weaker effect than in primary screening. From these 11 molecules, we resynthesized the 3 that maintained normalized reporter activation > 0.1 in retesting and found that 2 of them, Ribavirin and Liothyronine, elicited MutSβ-specific reporter activation across a range of concentrations (Figures S5K and S5L). However, similar to most strong hits from this collection (Table S8), these 2 molecules also activated our primary MutSα-specific reporter and thus did not show specificity for MutSβ-driven MMR.

Although much larger, the general collection produced only 5 hits (Figure 3G); however, among those hits, we identified an analog of cmpd1 and three novel molecules that share an identical chemical scaffold, differing at only two functional groups (R_1_ and R_2_; red inset in Figure 3G). Additionally, among compounds that scored just below our hit threshold, we identified a fourth compound with the same scaffold (normalized reporter activation = 0.07). We resynthesized these compounds and found that all four of them activated our MutSβ-specific reporter in a dose-dependent manner and, indicative of sensor specificity, did not activate our MutSα-specific reporter (Figure 4A). We refer to these compounds as our SBi series of molecules.

**Figure 4.**
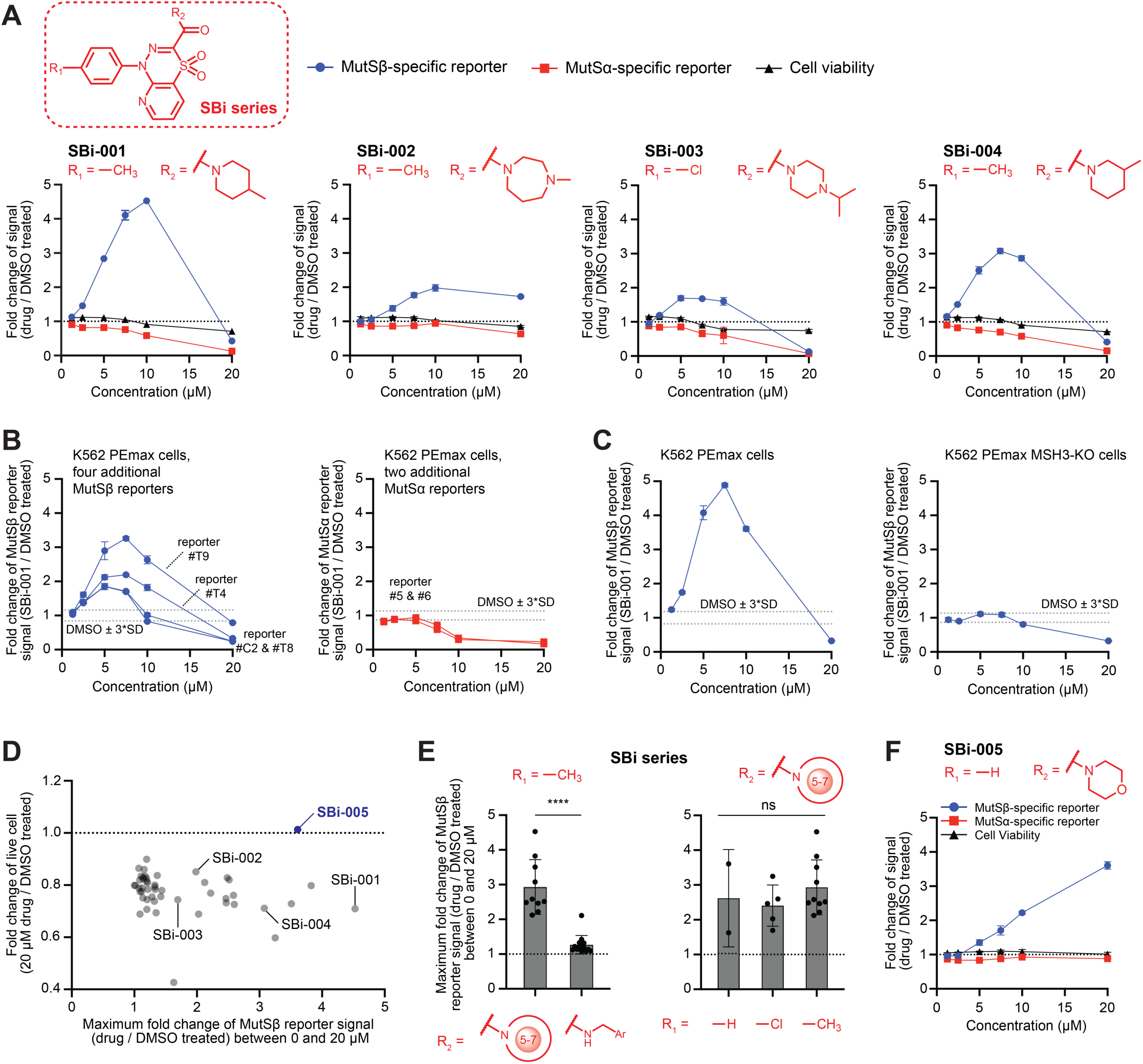
Cell-based validation and characterization of candidate MutSβ inhibitors. (A) Plots depict dose-dependent effects of SBi molecules on MutSβ-specific reporter activation (blue lines), MutSα-specific reporter activation (red lines), and cell viability (black lines, determined by Cell Titer-Glo) in K562 PEmax cells relative to the mean of DMSO controls. Dots represent fold changes in luminescence presented as means of n = 3 biological replicates and error bars indicate standard deviations. Core structure of SBi molecules included (top left). Relevant functional groups (R_1_ and R_2_) also included (plot insets). (B) Plots depict dose-dependent effects of SBi-001 on activation of additional MutSβ-specific reporters (left, blue lines) and additional MutSα-specific reporters (right, red lines) in K562 PEmax cells relative to the mean of DMSO controls. Dots represent fold changes in luminescence presented as means of n = 3 biological replicates, error bars indicate standard deviations, and dashed lines indicate 3 standard deviations from the mean of DMSO controls. (C) Plots depict dose-dependent effects of SBi-001 on MutSβ-specific reporter activation in K562 PEmax parental (left) or MSH3-KO (right) cells relative to the mean of DMSO controls. Dots represent fold changes in luminescence presented as means of n = 3 biological replicates, error bars indicate standard deviations, and dashed lines indicate 3 standard deviations from the mean of DMSO controls. (D) Testing SBi analogs for MutSβ-specific reporter activation (x-axis) and toxicity (y-axis). For activation of the MutSβ-specific reporter, each small molecule was tested at a range of concentrations (0 and 20 μM) and the maximal fold change relative to DMSO controls was reported. Since toxicity was monotonically related to drug concentrations, fold changes in live cells at 20 μM were reported (determined by Cell Titer-Glo). Dots represent means of n = 3 biological replicates. (E) Comparisons of SBi analogs with a methyl group as R_1_ and a different R_2_ (left) or with a nitrogen-containing 5-7 member ring as R_2_ and a different R_1_ (right) in K562 PEmax cells. Relevant functional groups (R_1_ and R_2_) are depicted (insets), wherein the 5-7 circle indicates a nitrogen-containing 5-7 member ring for which the other atoms on the ring can be substituted at any position. Each dot represents the maximal fold change in activation of the MutSβ-specific reporter relative to DMSO controls for one analog, selected from a range of concentrations, as in D. Bars represent means across analog groups and error bars indicate standard deviations. Mann-Whitney U test (left; ***, *p* < 0.0001) and Kruskal-Wallis test (right; ns, not significant, *p* >= 0.05) were used for statistical analysis. (F) Plot depicts dose-dependent effects of SBi-005 on MutSβ-specific reporter activation (blue line), MutSα-specific reporter activation (red line), and cell viability (black line, determined by Cell Titer-Glo) in K562 PEmax cells relative to the mean of DMSO controls. Dots represent fold changes in luminescence presented as means of n = 3 biological replicates and error bars indicate standard deviations. SBi-005 functional groups (R_1_ and R_2_) depicted (plot insets). Data in D-F are from the same experiment.

### Structure-activity relationship (SAR) analysis of candidate MutSβ inhibitors

In principle, hits from our screen could act through several indirect effects (e.g., nonspecific stimulation of prime editing or luciferase expression); however, because the SBi molecules identified in our screen activated our MutSβ-specific reporter but not our MutSα-specific reporter during counterscreening, those compounds are unlikely to operate through such effects. To confirm that selective reporter activation by these molecules was not limited to our primary reporter designs (e.g., did not represent selective suppression of only the exact DNA mispair specified in MutSβ-specific #T5), we next evaluated two of the molecules, SBi-001 and SBi-004, with several additional reporters, each designed to generate a distinct but still sensor-selective DNA mispair (Figures 2D–2G). Consistent with broad pathway-level inhibition of MutSβ-driven MMR, SBi-001 and SBi-004 activated four additional MutSβ-specific reporters, with similar dose-dependencies, and failed to activate two additional MutSα-specific reporters (Figures 4B and S6A). We therefore next asked if activation of our primary MutSβ-specific reporter by SBi-001 and SBi-004 was dependent on MutSβ, as would be expected for on-target pathway inhibition, and we found that neither compound activated that reporter in cells lacking MutSβ (Figures 4C and S6B). Altogether, these results suggest that SBi molecules selectively inhibit MutSβ-driven MMR.

Notably, at the highest concentration tested (20 μM), SBi compounds from our screen decreased signals from our MutSα/β-specific reporters relative to DMSO-treated controls (Figures 4A, 4B, and S6A). These effects, which mirrored loss of signals observed with cmpd1 (Figure S5A), coincided with reductions in cell viability and were thus likely due to nonspecific cellular toxicity (Figure 4A). To identify less toxic SBi compounds, we tested 45 analogs, each with the same or highly similar scaffolds and different functional groups (R_1_ and R_2_) (Figure 4D; Table S9). Comparative analysis showed that MutSβ-specific reporter activation was unaffected by small changes to R_1_ but strongly influenced by the amide amine at R_2_ (Figure 4E); that is, analogs with a 5-7 membered-ring tertiary amide at R_2_ exerted a larger effect than those with a secondary benzylic amide group. Altogether, 16 analogs (36%) reproduced activation of our primary MutSβ-specific reporter, and while none of these compounds produced a stronger effect than the most potent SBi molecule from our screen (SBi-001), one outperformed all other screen-nominated SBi molecules and showed no toxicity in K562 cells up to the highest concentration tested (20 μM; SBi-005) (Figures 4D, 4F and S6C). Active SBi compounds with less toxicity can thus be identified—albeit with possible cell type differences, as both SBi-001 and SBi-005 reduced reporter activation at the highest concentrations in RPE-1 hTERT cells (Figures S6D and S6E).

We next investigated changes to the SBi scaffold region (1H-pyrido[2,3-e][1,3,4]thiadiazine 4,4-dioxide). Across 53 analogs with scaffold modifications, none activated our MutSβ-specific reporter more strongly than SBi-001. However, several analogs demonstrated inverted reporter specificities, in that they elicited weak activation of our MutSα-specific reporter but showed no activation of the MutSβ-specific reporter (Figures S6F and S6G; Table S9). Scaffolds in these molecules often consisted of a benzene ring fused to a six-membered thiazine ring (4H-benzo[b][1,4]thiazine 1,1-dioxide) and thus contained two less nitrogen atoms than the fused pyridine and thiadiazine rings in the SBi scaffolds. We refer to such analogs as SAi molecules (Figures S6H-S6J). Intriguingly, we also found a single compound (Si-001) that weakly activated our MutSα-specific reporter and showed moderate MutSβ-specific reporter activity (Figures S6F and S6K). The scaffold in this molecule (1H-benzo[e][1,3,4]thiadiazine 4,4-dioxide) was similar to those in SBi and SAi compounds but had an intermediate number of nitrogen atoms: one less than in the SBi scaffold and one more than in the SAi scaffold. These results suggest that subtle changes to the bicyclic scaffold can alter MutS pathway sensitivity, possibly by differentially influencing interactions with MutSα and MutSβ.

### SBi compounds inhibit MutSβ binding to DNA mispairs

We next sought to define the molecular mechanisms by which our SBi compounds inhibit MutSβ-directed MMR. We first considered two indirect mechanisms: (1) reduction of MutSβ expression and (2) nonspecific interference of MutSβ binding to DNA from DNA intercalation. To determine whether SBi compounds affect MutSβ expression, we quantified MSH3 mRNA and protein levels in K562 PEmax cells with and without SBi-001 and SBi-005 (Figures S7A and S7B). Neither compound diminished expression at active concentrations. To evaluate the possibility of nonspecific DNA intercalation, we used surface plasmon resonance (SPR) and quantified the interaction of SBi-005 with a small DNA mispair (i.e., two looped-out CAGs within a 38 bp heteroduplex; Figure S7C). Unlike the known DNA intercalator mitoxantrone, which bound the mispaired DNA starting at low concentrations (>0.1 µM), we observed no evidence of an interaction with SBi-005 up to the highest concentration of compound tested (100 µM). These results suggest that SBi compounds do not inhibit MutSβ-directed MMR through these indirect mechanisms.

Given these results, we explored the possibility that SBi molecules act on MutSβ directly. We first asked whether SBi-005 alters binding of MutSβ to the same 38 bp (CAG)_2_ loop-out DNA using SPR and observed concentration-dependent reductions in normalized SPR signal, consistent with inhibition of MutSβ binding to the substrate (Figures 5A, S7D, and S7E; Methods). In parallel, we evaluated the impact of SBi-005 on MutSα using a heteroduplex harboring a canonical MutSα substrate (i.e., 38 bp DNA with a G-T mismatch), and observed a comparably modest effect, suggesting selectivity for MutSβ over MutSα (Figure 5B).

**Figure 5.**
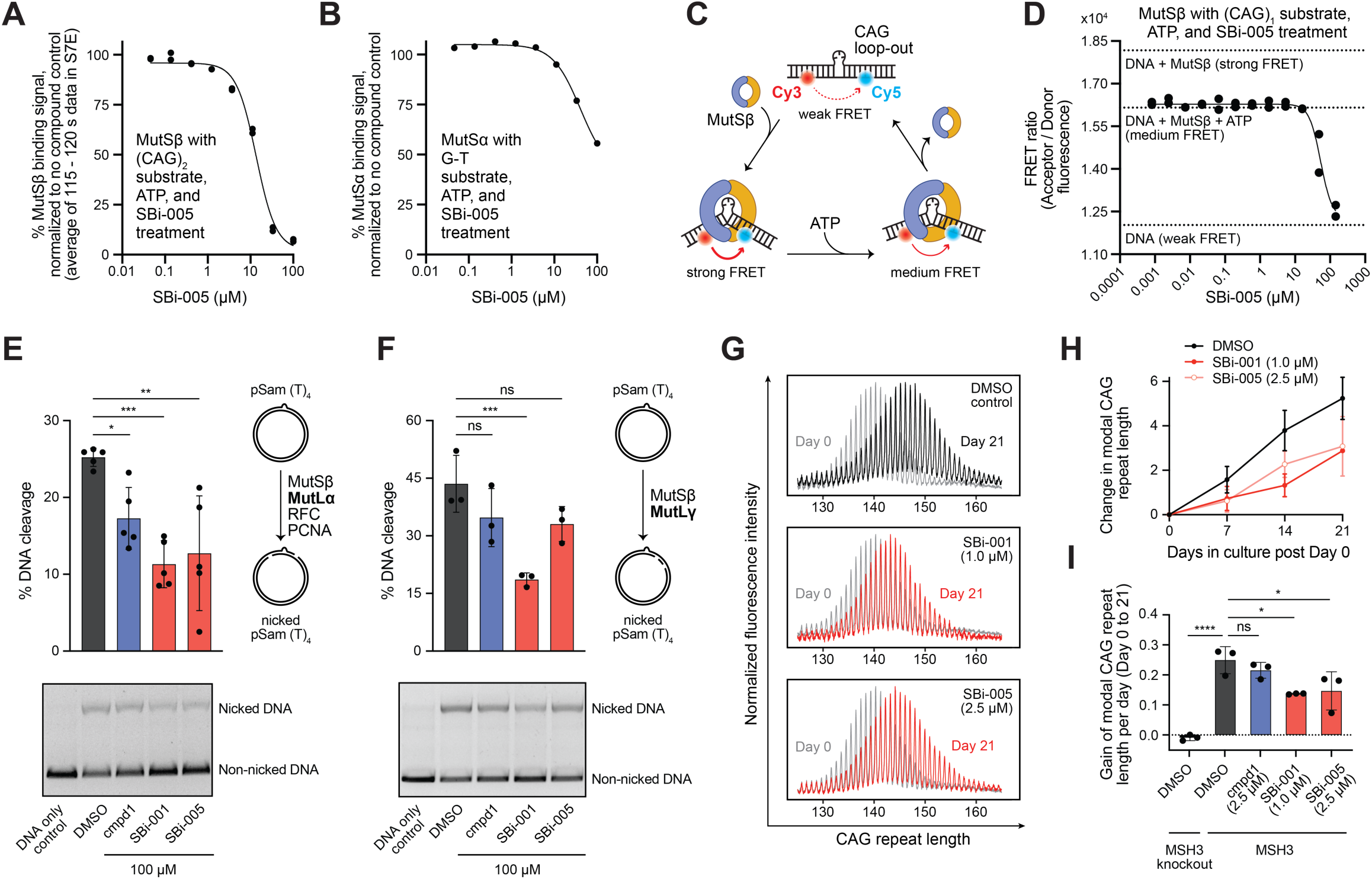
SBi molecules inhibit MutSβ binding to mispaired DNA and attenuate CAG expansion. (A) MutSβ binding to a 38 bp DNA containing a (CAG)_2_ loop-out at different concentrations of SBi-005 measured by SPR (as illustrated in Figure S7D). Dots represent means of mass response signals in resonance units (from 115 - 120 s, as in Figure S7E) from n = 2 biological replicates normalized to controls with no compound added. Black line represents a sigmoidal fit. (B) As in A but for MutSα binding to a 38 bp DNA containing a G-T mismatch. (C) Schematic of FRET assay. Three cycling states from reactions performed in the presence of ATP are depicted: MutSβ-unbound DNA (top, weak FRET), MutSβ-bound inducing DNA bending (bottom left, strong FRET), and ATP-dependent MutSβ sliding along and/or falling off from DNA (bottom right, medium FRET). When both MutSβ and ATP are present, cycling among the three states can produce medium FRET. (D) MutSβ binding to a 70 bp DNA containing a (CAG)_1_ loop-out at different concentrations of SBi-005 measured in the presence of ATP by FRET assay. Dots represent FRET ratios from n = 2 biological replicates and the solid black line represents a sigmoidal fit. Control signals measured without compound treatment are indicated by black dashed lines. (E) *In vitro* cleavage of a plasmid containing a (T)_4_ loop-out by recombinant MutSβ-MutLα in the presence of DMSO, cmpd1, SBi-001, or SBi-005. Dots in barplot (upper left) represent percentages of cleavage, bars indicate means of n = 5 biological replicates, and error bars indicate standard deviations. Ordinary one-way Analysis of Variance (ANOVA) was used for statistical analysis followed by Dunnett’s multiple comparisons test (***, adjusted *p* value < 0.001; **, adjusted *p* value < 0.01; *, adjusted *p* value < 0.05). A representative gel image (lower left) and a schematic of the experiment (upper right) are also included. (F) *In vitro* cleavage of a plasmid containing a (T)_4_ loop-out by recombinant MutSβ-MutLγ in the presence of DMSO, cmpd1, SBi-001, or SBi-005. Dots in barplot (upper left) represent percentages of cleavage, bars indicate means of n = 3 biological replicates, and error bars indicate standard deviations. Ordinary one-way ANOVA was used for statistical analysis followed by Dunnett’s multiple comparisons test (***, adjusted *p* value < 0.001; ns, not significant, adjusted *p* value >= 0.05). A representative gel image (lower left) and a schematic of the experiment (upper right) are also included. (G) Plots depict representative distributions of CAG repeat lengths from a cellular model of repeat expansion (i.e., RPE-1 hTERT cells with exogenous mHTT exon 1) at initial sampling (defined as Day 0, grey, two days post treatment with DMSO, SBi-001, or SBi-005) and 21 days later. Modal CAG repeat lengths were ∼130-140 at initial sampling. (H) Changes in modal CAG repeat length observed in a cellular model of repeat expansion after 21 days of growth in the presence of DMSO, SBi-001, and SBi-005 since Day 0 (vehicle and compounds replenished every 3-4 days). Dots represent means of n = 3 biological replicates and error bars indicate standard deviations. (I) Per day changes in modal CAG repeat length observed in a cellular model of repeat expansion after 21 days of growth in the presence of DMSO, SBi-001, and SBi-005 since Day 0 (vehicle and compounds replenished every 3-4 days). Dots represent per day averages, bars indicate means of n = 3 biological replicates, and error bars indicate standard deviations. Ordinary one-way ANOVA was used for statistical analysis followed by Tukey’s multiple comparisons test (****, adjusted *p* value < 0.0001; *, adjusted *p* value < 0.05; ns, not significant, adjusted *p* value >= 0.05). Data in G-I are from the same experiment.

We next deployed an orthogonal MutSβ binding assay, which uses Förster resonance energy transfer (FRET) to measure heteroduplex bending upon interaction with MutSβ (herein, a 70 bp substrate with one CAG loop-out; Figure 5C). When performed with ATP, this assay reports an equilibrium FRET ratio that reflects the distribution of three cycling reaction states: MutSβ bound to the loop-out, ATP-dependent sliding of MutSβ on DNA, and unbound MutSβ.^81^ Further supporting compound-dependent inhibition of MutSβ-DNA binding, SBi-005 caused dose-dependent reductions in this equilibrium ratio, with higher SBi-005 concentrations producing ratios closer to controls performed without MutSβ (Figure 5D). Notably, in the absence of ATP, SBi-005 exhibited lower potency (Figure S7F), possibly due to a higher threshold for inhibition in that context caused by formation of more stable MutSβ-DNA complexes, as expected from reduced release of MutSβ from DNA without ATP, and as evident in stronger MutSβ-induced FRET ratios.

Downstream of MutSβ, MutL heterodimers (MutLα or MutLγ) nick the mispaired DNA to coordinate repair. We therefore next asked whether SBi molecules also inhibit these events. In an established *in vitro* assay,^69^ SBi-001 and SBi-005 compounds both compromised MutSβ:MutLα-mediated cleavage of a plasmid harboring an extrahelical (T)_4_ loop-out, notably with stronger effect than cpmd1, which we included as a positive control (Figure 5E). Additionally, in a similar assay, SBi-001 inhibited MutSβ:MutLγ-mediated cleavage of the same substrate; although neither cpmd1 nor SBi-005 elicited a similar effect in this assay, possibly due to differences in the potencies required for inhibition (Figure 5F). Given these results, we next tested the effect of SBi-001 on a separate nuclease, FAN1, and observed no difference in cleavage with or without the compound, indicating that SBi compounds do not indiscriminately inhibit endonuclease activities (Figure S7G). Alongside our other findings, these results support a model wherein SBi compounds inhibit MutSβ-driven MMR by interfering with MutSβ substrate recognition, upstream of DNA cleavage.

### SBi compounds attenuate CAG repeat expansion

Identifying selective inhibitors of MutSβ-directed MMR is an important step; however, therapeutic potential requires that inhibition from candidate compounds translates into suppression of repeat expansion. We therefore asked whether our SBi molecules could attenuate repeat expansion in RPE-1 hTERT cells harboring a stably integrated (CAG)_129_ mHTT exon 1. Cells were grown with SBi-001, SBi-005, and cmpd1 at 1 µM or 2.5 µM, depending on expected toxicities, collected for initial analysis after 2 days (designated day 0 samples), and then grown for 21 days with compound replenishment and intermittent sampling. CAG tracts were amplified from genomic DNA and analyzed by capillary electrophoresis (Figure 5G). Comparison of modal repeat lengths revealed significantly decreased (CAG)_n_ expansion in cells treated with SBi-001 and SBi-005, compared to DMSO controls (Figures 5H and 5I). Moreover, despite potential for attenuation of repeat expansion to arise from indirect effects on the cell cycle (i.e., reduced occupancy in S phase resulting in fewer replication-associated strand slippages), cell cycle analysis of similarly treated RPE-1 hTERT (CAG)_129_ cells showed only mild, inconsistent changes, suggesting that observed effects were more likely caused by inhibition of MutSβ (Figure S7H). These results solidify the utility of our prime editing-based reporter assays for identifying MutSβ inhibitors relevant to cellular repeat expansion.

## DISCUSSION

Previous studies have suggested that prime editing forms transient DNA mispairs at targeted genomic loci,^51–55^ and because prime editing is easily programmable, the technology allows flexible modeling of those editing intermediates directly in genomes (e.g., base-base mismatches and insertion/deletion loop-outs). We exploited the scalability of prime editing to profile the effects of core MMR genes across thousands of specified mispairs. Our results revealed information-rich “fingerprints” of MMR gene functions, resolved the substrate preferences of MutSα and MutSβ at scale, and isolated poorly characterized MMR activities in cells. Among key observations, we found that *MLH3* suppresses small deletions and promotes similarly sized insertion edits. These phenotypes, alongside those observed in cells lacking *MSH3*, support a mechanism of loop-directed MMR, which has been defined biochemically^69,70^ but not resolved in cells.

This noncanonical, MutLγ-mediated mechanism favors retention of looped-out DNA in repaired duplexes and is thus highly relevant to repeat expansion. We identified phenotypes consistent with activity of this mechanism in cells and thus provide a straightforward way to investigate it further (e.g., characterize sequence features).

While many questions remain about MutLγ-mediated MMR, less is known about the role of MutLβ in DNA repair. Like MutSβ, MutLβ contributes to (CAG)_n_ expansion^8,9,18,19^ and is therefore an HD drug target of interest^82^; however, because MutLβ lacks the DQHA(X)_2_E(X)_4_E endonuclease motif present in other MutL complexes^67^ and exhibits limited MMR function *in vitro*,^83^ its role in MMR has been unclear. Addressing this gap, we found that loss of MutLβ in PMS1-KO cells produced phenotypes similar to those caused by loss of MutLγ in MLH3-KO cells; that is, we observed nearly identical changes in deletion editing and more subtle but concordant effects on insertions (Figures 1F, 1I, and S2D). These similarities suggest that, like MutLγ, MutLβ promotes loop-directed MMR. Supporting this idea, a recent study showed that MutLβ enhances MutSβ-stimulated MutLγ endonuclease activity *in vitro* and blocks an inhibitory, potentially antagonistic effect from MutSα.^84^

Two additional, related observations from our genetic screens are that (1) MutLα deficiency in PMS2-KO cells produced phenotypes similar to those from MLH1-KO cells, which lack all three MutL complexes, and (2) phenotypes attributed to *MLH1* did not generally account for effects caused by targeted disruption of MutLβ or MutLγ in PMS1-KO and MLH3-KO cells, respectively. These observations suggest that epistatic relationships exist among MutL complexes, and given that MutLβ and MutLγ are promising HD drug targets^85^ but MutLα is not, resolving these relationships will be of high interest. Our genetic screening framework, which should pair easily with technologies that disrupt multiple genes,^86–94^ will enable such future studies.

In addition to providing insight into cellular mechanisms of MMR, our genetic screens resolved classes of prime edits selectively suppressed by the two MMR sensors, MutSα and MutSβ. Guided by these results, we built a set of scalable, sensor-selective MMR reporters with several advantages over alternative methods for evaluating MMR in cells. Unlike other methods, these reporters measure separable MMR activities within only a few days, have high dynamic range, and, due to having a positive readout (i.e., luciferase expression), minimize false positives from cellular toxicity. In proof-of-principle studies, we used these reporters to identify and characterize a series of selective MutSβ inhibitors. Additional studies will be required to determine the therapeutic potential of these and other SBi-type molecules; however, those identified herein already provide valuable tools for investigating MutSβ function and for exploring MutSβ targeting strategies (e.g., identifying druggable pockets). More broadly, the identification of these molecules definitively establishes our reporters as useful tools for therapeutic discovery. Applied across larger libraries, we expect these reporters to identify even more potent and less toxic selective inhibitors of MutSβ, and because our MutSβ-specific reporters register loss of multiple HD modifier genes (i.e., *MLH3* and *PMS1*), such screens may also uncover compounds against those targets.

Notably, our reporters can also be applied *post hoc* to characterize MMR inhibitors found through orthogonal approaches (e.g., from *in vitro* screens), as we showed for cmpd1 and one PMS2 inhibitor. Moreover, as drug development efforts advance, these reporters can be used to compare pharmacological inhibition of MMR to loss of MMR components or expression of inactive variants, each of which may block MMR via distinct mechanisms. Additionally, with straightforward modifications, our reporters could be adapted to enable massively parallel characterization of MMR genetic variants, including those from genome-wide association studies, or to catalog regulators of separable MMR pathways (e.g., loop-directed MMR) with large-scale genetic screens. Such applications will primarily require a simple design change; that is, the current luciferase-encoding reporter gene would need to be replaced by a gene more amenable to pooled screening (e.g., GFP).^65^

Herein, we also establish new ways to improve prime editing. To date, multiple studies have confirmed that canonical MMR suppresses prime editing with small deletions, small insertions, and substitutions.^51–57,65,95^ To overcome this limitation, methods for transiently inhibiting MMR have been developed; however, these methods rely on expression of a dominant negative protein^51^ or an engineered peptide^95^ and thus are limited by delivery challenges. Compared to these methods, the small molecule inhibitors identified and characterized in our study offer advantages, including easy and flexible delivery. For improvement across a wide range of edit types, we recommend the PMS2 inhibitor. Additionally, our findings establish an expanded model of prime editing, wherein installation of small insertions, but not deletions, is aided by an endogenous MMR subpathway. This finding changes our understanding of prime editing and explains why small insertions are generally preferable to deletions in applications of editing in MMR-proficient cells.

Altogether, our work establishes several versatile, cell-based assays for interrogating MMR. These systems provide powerful tools for studying mechanisms of genome maintenance and somatic instability in repeat expansion diseases, with demonstrated capacity to identify inhibitors of significant therapeutic value.

## LIMITATIONS OF THE STUDY

Several limitations of our study should be considered. With respect to assay development, we highlight two limitations that complicate MMR pathway modeling: (1) While programmed heteroduplex structures generated by prime editing likely arise as expected, unprogrammed structures may also form (e.g., driven by alternative flap hybridizations). Given this potential, we focused on median phenotypes when analyzing results from our genetic screens (i.e., phenotypes observed across edit classes), rather than individual effects; however, by accounting for heteroduplex ensembles, future studies may further resolve MMR subpathways. (2) Because prime editing-induced heteroduplex structures invariably form near a DNA strand discontinuity, programmed mispairs recognized by MutSα and MutSβ will likely always stimulate canonical, MutLα-catalyzed repair. This feature unavoidably complicates isolation of other MMR processes, especially those that may operate on continuous, unbroken DNA. Moreover, because edit-associated strand discontinuities occur invariably on the 3′ side of programmed mispairs, repair though canonical MMR may be biased towards 3′ nick-directed mechanisms, rather than 5′ nick-directed ones.^96^ As highlighted above, future studies may address these challenges by performing experiments in defined genetic backgrounds, for example by knocking out *PMS2* to avoid MutLα activity or overexpressing DNA ligase to promote ligation of the 3′ nick. Additionally, deploying our assays in non-dividing cells (e.g., cultured or iPSC-derived neurons) may further improve translational relevance for specific applications.

With respect to the early-stage MutSβ inhibitors identified in this study, we note generally pervasive toxicity. Two lines of evidence argue that this toxicity is not driven by on-target MutSβ inhibition: (1) our MSH3-KO cells, which lack MutSβ, were comparably viable, with no obvious growth defects, and (2) one analog of these molecules (SBi-005) retained targeted activity but showed significantly less toxicity in K562 cells, suggesting a more tolerable chemical structure, at least for that cell type. Additional studies will be required to distinguish off-target liabilities and determine the precise mechanisms of action of these molecules, however, pending such studies, this consideration motivates continued search for additional small molecules suitable for therapeutic development—efforts that will be greatly accelerated by our advances.

## Acknowledgements

We thank Aaron Lin and other Adamson Lab members for discussions and suggestions. We thank Wei Wang, Jennifer M. Miller, Jean Arly Volmar, Julia E. Good, and Alexander Harms from the Genomics Core Facility at Princeton University, and Christina DeCoste and Jailene Garcia from the Flow Cytometry Resource Facility at Princeton University (partially supported by the Rutgers Cancer Institute of New Jersey, NCI-CCSG P30CA072720-5921). We thank Brinda Prasad and Simon Noble at the CHDI Foundation for their advice and suggestions on the manuscript, and Kezhi Chen at the University of Chicago for chemistry discussions. This work was supported by the nonprofit CHDI Foundation Inc. CHDI Foundation is a nonprofit biomedical research organization exclusively dedicated to collaboratively enabling the development of therapeutics that substantially improve the lives of those affected by Huntington’s disease. Research in the Adamson lab was also supported by the National Institutes of Health (NIH) under award numbers R35GM138167 and RM1HG009490, Ludwig Cancer Research, and Princeton University, in part through the Intellectual Property Accelerator Fund. Research in the Balmus Lab was also supported by the UK Dementia Research Institute, with funding from the UK MRC, and the Romanian Ministry of Research, Innovation, and Digitization (grant #PNRR-III-C9-2022-I8-66, contract 760114). Yangwode Jing was supported by the George B. Rathmann *51 Graduate Fellowship and the James Mi *91 and Teresa Mi S *91 Graduate Fellowship from Princeton University. Jun Yan was supported by a fellowship provided by the China Scholarship Council (CSC), based on the April 2015 Memorandum of Understanding between the CSC and Princeton University. Yangwode Jing dedicates this work to his grandparents and parents, by whom he was raised up.

## Author contributions

Conceptualization, Y.J., J.Y., and B.A. Generation and characterization of cell lines, Y.J. with J.Y. and P.R. Self-targeting prime editing screens, Y.J. with J.Y. and J.L. Formal analysis of prime editing screens, Y.C. with Y.J. and J.Y. Interpretation of prime editing screens, Y.J., J.A.H., and B.A. Construction of MutSα- and MutSβ-specific reporters, Y.J. with J.Y. and K.W. Small molecule screen, Y.J. with J.Y. and H.K. SAR studies, Y.J., H.K., L.L., and M.F. SPR and FRET experiments and analysis, H.D., V.D., and M.T. *In vitro* MMR assays, I.S. Repeat expansion assays, D.Y., J.W., and K.H. All other experiments and data analysis: Y.J. Project administration, M.F., T.F.V., and B.A. Supervision, Y.P., G.B., P.C., M.F., and B.A. Writing, Y.J. and B.A. with the input from all authors.

## EXPERIMENTAL MODEL AND SUBJECT DETAILS

K562 PEmax parental cells and MLH1-KO cells were previously described.^56,65^ Lenti-X 293T cells (632180) were from Takara. HeLa (CCL-2), U-2 OS (HTB-96), HCT 116 (CCL-247), and RPE-1 hTERT (CRL-4000) were from ATCC. HAP1 (parental, C631; MSH3-KO, HZGHC000656c001; MSH6-KO, HZGHC000653c003; MSH2-KO, HZGHC024799c006; MLH1-KO, HZGHC000343c021) were from Horizon. K562 PEmax cells were cultured in RPMI 1640 medium, HEPES (Gibco, 22400089) supplemented with 10% (v/v) fetal bovine serum (FBS; Gibco, A52567-01) and 1x Penicillin-Streptomycin-Glutamine (Gibco, 10378-016), also referred to as “complete RPMI medium”. K562 PEmax cells were grown between 4e5 to 1e6 cells / mL. Lenti-X 293T and HeLa cells were cultured in Dulbecco’s modified Eagle’s medium (Corning, 10-013-CV) supplemented with 10% (v/v) FBS (Gibco, A52567-01) and 1x Penicillin- Streptomycin (Corning, 30-002-Cl). HAP1 cells were cultured in Iscove’s Modified Dulbecco’s medium (IMDM; Cytiva HyClone^TM^, SH30228.01) supplemented with 10% (v/v) FBS (Gibco, A52567-01) and 1x Penicillin-Streptomycin (Corning, 30-002-Cl). HCT-116 and U-2 OS cells were cultured in McCoy’s 5A (Modified) Medium (Gibco, 16600-082) supplemented with 10% (v/v) FBS (Gibco, A52567-01) and 1x Penicillin-Streptomycin (Corning, 30-002-Cl). RPE-1 hTERT cells were cultured in DMEM/F12 medium with L-Glutamine and 15 mM HEPES (Gibco, 11330-032) supplemented with 10% (v/v) FBS (Gibco, A52567-01), 1x Penicillin-Streptomycin (Corning, 30-002-Cl), and 10 μg/mL hygromycin B (Goldbio, H270-2.5), with the following exception: In select experiments (Figures 5G-5I and S7H), we used a second RPE-1 hTERT cell line, called RPE-1 hTERT (CAG)_129_, details of which can be found below (*Cell-based CAG repeat expansion assay*). For experiments herein, this cell line was cultured in Dulbecco-modified essential medium (DMEM), high glucose (Gibco, 11965092), supplemented with 10% (v/v) FBS (Gibco, A52567-01), 1% (v/v) MEM NEAA (Gibco, 11140050), and 1x Penicillin-Streptomycin-Glutamine (Gibco, 10378-016). All cells used in this study were cultured at 37 °C in a humidified incubator with 5% CO_2_ (v/v). Cell lines were authenticated by either suppliers or short tandem repeat profiling by ATCC and routinely tested negative for mycoplasma.

Small molecules used in this study include cmpd1 (purchased from Ambeed, A1110529), PMS2i (synthesized by WuXi Apptec Co., Ltd.), Ribavirin (purchased from MedChemExpress, HY-B0434), Levocarnitine propionate hydrochloride (purchased from MedChemExpress, HY-B0932), and Liothyronine (purchased from MedChemExpress, HY-A0070A). Other small molecules were ordered from ChemDiv Inc. Prior to cell-based experiments, small molecules were dissolved in dimethyl sulfoxide (DMSO, EMD Millipore, 317275-500ML). A summary of compounds used in this study can be found in Table S9.

## METHOD DETAILS

### Expression plasmids

Plasmids for transfection or electroporation of pegRNAs were cloned using pU6-pegRNA-GG-acceptor (Addgene, 132777). Briefly, pegRNA sequences (affixed to a modified mouse U6 promoter) were PCR amplified from gene fragments (Twist Bioscience), digested with BamHI-HF and XhoI, purified, and then assembled into BamHI-HF- and XhoI-digested backbone. Plasmids for transfection or electroporation of *MSH3*, *MSH6*, and *MSH2* were ordered from GenScript (OHu20501D, OHu25314D, and OHu20167D in pcDNA3.1-C-(k)DYK). In these initial plasmids, MMR cDNAs were tagged with FLAG (DYKDDDK, C-terminal). We removed this tag from each vector, creating modified plasmids for expression of untagged MSH3, MSH6, and MSH2 (pYJ067, pYJ080, and pYJ189, respectively). Expression of MMR genes delivered by transfection or electroporation was driven by the CMV promoter. Finally, plasmids for expression of MMR proteins via lentiviral transduction were assembled using cDNAs from pYJ067, pYJ080, pYJ189 or plasmids containing MutL subunits ordered from Twist Bioscience and FUGW lentiviral backbone (Addgene, #14883). For expression of PE7 proteins via lentiviral transduction, PE7 were PCR amplified from pCMV-PE7 (Addgene, #214812) and assembled with a pU6-sgRNA EF1Alpha-puro-T2A-BFP (Addgene #60955)-derived lentiviral backbone, IRES sequences (Addgene, #210590), and miRFP680 encoding sequences.^97^ All plasmids were prepared with NucleoSpin Plasmid, Mini kit (Macherey-Nagel, 740588.250), ZymoPURE II Plasmid Midiprep Kit (Zymo Research, D4201), or ZymoPURE II Plasmid Maxiprep kit (Zymo Research, D4203).

### Backbone plasmids for self-targeting libraries

Throughout this study, we used the PE2 approach without the nicking guide RNA.^46^ We generated two plasmids to enable our self-targeting prime editing screens. These plasmids, which served as backbones for library cloning (*Cloning of HEK3 (lYJ003) and FANCF (lYJ004) self-targeting libraries*), have several key components: (1) a pegRNA expression cassette composed of a modified mouse U6 promoter, spacer for HEK3 or FANCF (5′-GGCCCAGACTGAGCACGTGA-3′ for HEK3 and 5′-GGAATCCCTTCTGCAGCACC-3′ for FANCF), and an SpCas9 guide RNA scaffold sequence (5′-GTTTTAGAGCTAGAAATAGCAAGTTAAAATAAGGCTAGTCCGTTATCAACTTGAAAAAGTGGCACCGAGTCGGTGC-3′);^98^ (2) a cassette designed to enable subsequent library construction composed of an Esp3I recognition sequence, mRFP1, a SbfI recognition sequence, an XhoI recognition sequence, the reverse complement of the target sequence from endogenous HEK3 or *FANCF*, where the pegRNAs can bind and install the edits, and an i7 annealing region (5′-AGATCGGAAGAGCACACGTCTGAACTCCAGTCAC-3′); and (3) a selection marker cassette composed of an EF1a promoter, puromycin resistance gene (PuroR), T2A, and TagBFP.

To build these two plasmids, we modified an existing pegRNA expression plasmid (pAC025, Addgene, 227597).^56^ Notably, inclusion of mRFP1 in the second cassette enables differentiation of colonies with inserts that replace sequences between the Esp3I and XhoI recognition sequences (white colonies) from those without (red colonies) during library construction.

### Design of HEK3 (lYJ003) and FANCF (lYJ004) self-targeting libraries

We constructed two self-targeting libraries (lYJ003 and lYJ004) for our prime editing screens (Table S1). These libraries contained the following key elements (5′ to 3′; Figures 1C, S1A, and S1B): (1) a pegRNA expression cassette driven by a modified mouse U6 promoter, for expression of individual pegRNAs with a single spacer sequence, standard constant region of SpCas9 guide RNA scaffold,^98^ and variable extension sequences (6000 per library), and (2) a single target sequence (90 nt of HEK3 for lYJ003 and *FANCF* for lYJ004) adjacent to a 12-mer barcode (generated by an R package called DNABarcodes^99^). Each of these libraries specified 6000 edits: 2539 (HEK3) or 2542 (FANCF) insertions, 3263 (HEK3) or 3283 (FANCF) substitutions, and 96 (HEK3) or 97 (FANCF) deletions, and had 102 (HEK3) or 78 (FANCF) negative controls. Across the two libraries, edits had the same general patterning:

#### Insertions

In each library, we included insertions ranging from 1 bp to 25 bp with starting positions programmed directly adjacent to the predicted nick made by PEmax (+1 position) or six base pairs away (+7). All library insertions were designed from 30 “parent” sequences, containing 26 standard utility sequences (e.g., FLAG tag) and 4 repeat sequences related to human disease such as (CAG)_n_. To generate a diverse but related set of insertions, we included each “parent” sequence, their reverse complement, and successive truncations thereof (i.e., full length sequence minus 1 from the 3′ end repeatedly until only 1 nt from the 5′ end remained) into each library in each start position. Then, to keep library elements unique, we removed any repeated insertions. All insertions, within each of the two libraries, were designed with the same primer binding site sequences (PBS, 5′-CGTGCTCAGTCTG for HEK3 and 5′-GCTGCAGAAGGGAT for FANCF) and with reverse transcription templates (RTT) containing the same homology regions (5′-TGGAGGAAGCAGGGCTTCCTTTCCTCTGCCATCA-3′ for HEK3 +1 position, 5′-TGGAGGAAGCAGGGCTTCCTTTCCTCTG-3′ for HEK3 +7 position, 5′-GAGACCGCCAGAAGCTCGGAAAAGCGATCCAGGT-3′ for FANCF +1 position, 5′-GAGACCGCCAGAAGCTCGGAAAAGCGAT-3′ for FANCF +7 position).

#### Substitutions

We included substitutions of four types in each library: (1) All possible consecutive substitutions of 1-5 bp, starting directly adjacent to the predicted PEmax nick (+1 position) or two (+3) or four (+5) positions away. For each start site, these substitutions include 3 edits of 1 bp, 9 edits of 2 bp, 27 edits of 3 bp, 81 edits of 4 bp, and 243 edits of 5 bp for a total of 363 edits per start site and 1089 total edits; (2) All possible consecutive substitutions of 6 bp, starting at position +1 for a total of 729 such edits; (3) A select set of randomly generated consecutive substitutions ranging in length from 7-25 bp. These edits include 50 substitutions per length starting at position +1 for a total of 950 such edits; (4) A select set of non-consecutive substitutions, wherein the first substitution is at position +1 and the last is 4 or 5 bp away. These edits include 306 “complex substitutions” of 5 bp and 243 of 6 bp, respectively, for a total of 549 edits. Here too, we removed any repeated edits.

#### Deletions

In each library, we included deletions ranging from 1 bp to 25 bp with starting positions programmed directly adjacent to the predicted PEmax nick (+1 position) or two (+3) or four (+5) or six (+7) base pairs away. These deletions include 1 edit of each length and 25 edits per start site for a total of 100 such edits. Here too, we removed any repeat edits.

#### Negative controls

We also included two sets of negative controls in each library: (1) “non-annealing” pegRNAs containing scrambled extensions that should not anneal to the target site and (2) “no edit” pegRNAs with extensions that perfectly match the target site.

### Cloning of HEK3 (lYJ003) and FANCF (lYJ004) self-targeting libraries

To construct our self-targeting libraries, pegRNA extension sequences for either HEK3 or *FANCF* containing edits described above were ordered as oligonucleotide pools (Twist Bioscience) and cloned into one of two plasmids (*Backbone plasmids for self-targeting libraries*). Oligonucleotides within these pools contained the following elements (5′ to 3′): an annealing region for PCR amplification, an Esp3I recognition site, the pegRNA extension, a 7-mer polyT (transcription termination signal), a BamHI recognition site, an SbfI recognition site, a 12-mer barcode that uniquely corresponds to the paired pegRNA extension, a XhoI site, and another annealing region for PCR amplification. For cloning purposes, extensions and barcodes with Esp3I, BamHI, SbfI, or XhoI recognition sites were removed prior to ordering.

Cloning of each library was performed separately in several steps: (1) Sequences within each oligonucleotide pool were amplified by PCR using a pair of primers covalently labelled with biotin on their 5’-ends (oYJ379 and oYJ380; Table S4). PCR mixtures were assembled with NEBNext Ultra II Q5 Master Mix (M0544L), and reactions were performed at 98 °C for 30 s, then 12 cycles of [98 °C for 10 s, 62 °C for 20 s, and 72 °C for 30 s], followed by 72 °C for 2 min; (2) Amplified oligonucleotide pools were then purified using Oligo Clean & Concentrator (Zymo, D4060), according to manufacturer’s instructions, and sequentially digested with FastDigest XhoI and Esp3I (Thermo Scientific, FD0694 and FD0454); (3) Digested adaptors (with biotin) were removed using Dynabeads™ MyOne™ Streptavidin T1 from the Dynabeads™ Streptavidin Trial Kit (Invitrogen, 65801D) and remaining products (“inserts”) were purified with Oligo Clean & Concentrator; (4) Purified inserts were next ligated into corresponding backbones linearized by sequential digestion with FastDigest XhoI and Esp3I (Thermo Scientific, FD0694 and FD0454) and purified with Zymoclean Gel DNA Recovery Kit (Zymo, D4001). Ligation mixtures were assembled with ∼150 ng of backbone, ∼38 ng inserts, 1.5 µL T4 DNA ligase (NEB, M0202T), and water to 20 µL, and reactions were performed at 16 °C for 16 hrs; (5) Ligated product was then purified using SPRIselect reagent (Beckman Coulter, B23318) with a 0.7X left side selection following manufacturer’s instructions and electroporated into Endura DUOs electrocompetent cells (Biosearch Technologies, 60242-2) using a Gene Pulser Xcell electroporation system (BIO-RAD, 1652660); (6) After recovery at 37 °C, bacteria were grown in 200 mL LB broth supplemented with 70 µg/mL carbenicillin for 24 hours at 30 °C; (7) Finally, plasmid libraries were extracted using ZymoPURE II Plasmid Maxiprep Kit (Zymo, D4203).

To assess library quality, pegRNA cassettes and target regions were amplified and sequenced. Amplification was performed with a flanking 5′ primer (5′-AATGATACGGCGACCACCGAGATCTACACGCACAAAAGGAAACTCACCCT-3′) and 3′ indexing primer (5′-CAAGCAGAAGACGGCATACGAGATNNNNNNNNGTGACTGGAGTTCAGACGTGTGCTCTTC-3′) using the following program: 98 °C for 30 s, then 12 cycles of [98 °C for 10 s, 62 °C for 20 s, and 72 °C for 30 s], followed by 72 °C for 2 min. PCR products were purified using SPRIselect, quantified by a Qubit 4 Fluorometer, analyzed with an Agilent 2100 Bioanalyzer, and sequenced using a NovaSeq 6000 SP Reagent Kit v1.5 (300 cycles; Illumina, 20028400) with R1 = 190 nt (using oSL055, 5′-GTGTGTTTTGAGACTATAAGTATCCCTTGGAGAACCACCTTGTTG-3′ as a custom sequencing primer), I1 = 8 nt i7 index, and R2 = 135 nt. The libraries were well-balanced (Figure S1C).

### Lentivirus packaging and transduction

Lentivirus was prepared and cells were transduced using standard protocols. Materials for preparation were Lenti-X 293T cells, transfer plasmid(s), a packaging plasmid mix containing pALD-Rev-A, pALD-GagPol-A, and pALD-VSV-G-A (Aldevron), TransIT-LT1 (Mirus, MIR 2300), Opti-MEM I Reduced Serum Medium (Gibco, 31985070), and ViralBoost reagent (ALSTEM, VB100). Virus-containing supernatant was typically stored at -80 °C. Prior to transduction, all cells were isolated by centrifugation and resuspended into media with 6-8 μg / mL polybrene (Santa Cruz Biotechnology, sc-134220). K562 PEmax cells were transduced with centrifugation (“spinfection” at 1000 x g for 2 hours at room temperature in 6-well plates), and after centrifugation, cells were resuspended by swirling. Cells were incubated with virus for at least 48 hours before passaging or analysis. Transductions performed for prime editing screens are described with additional details below (*Self-targeting prime editing screens*).

### Cloning MutSα/β-specific reporter constructs

Reporter plasmid sequences are included in Table S7. Our MutSα/β-specific reporters have two key components: (1) a pegRNA expression cassette and (2) a reporter gene cassette that can be induced by prime editing (Figures 2D and 2E). To build the pegRNA expression cassette, we assembled a modified mouse U6 promoter (gene fragment, Twist Biosciences), spacer sequence, modified SpCas9 guide RNA scaffold (5′-GTTTAAGAGCTATGCTGGAAACAGCATAGCAAGTTTAAATAAGGCTAGTCCGTTATCAACTTGAAAAA GTGGCACCGAGTCGGTGC-3′),^100^ relevant extension sequence, trimmed evopreq1 riboswitch (5′-CGCGGTTCTATCTAGTTACGCGTTAAACCAACTAGAA-3′, only for MutSα reporters),^101^ and polyT transcription termination signal (5′ to 3′). To build the reporter gene cassette, we assembled an SFFV promoter, the corresponding pegRNA target site, the NanoLuc luciferase gene^102^ from pTRE3G-NlucP (Addgene, 162595), an IRES2 from pLenti-DsRed_IRES_eGFP (Addgene, 92194), and mCherry from pHR-SFFV-KRAB-dCas9-P2A-mCherry (Addgene, 60954) (5′ to 3′). We cloned these reporters in two steps: First, we inserted all elements except the pegRNA sequence, SFFV promoter, and corresponding pegRNA target into a lentiviral backbone using NEBuilder HiFi DNA assembly. The resulting plasmid contained two BsmBI restriction digestion sites between the mU6 promoter and luciferase gene. Second, we inserted pegRNA sequences, SFFV promoter, and targets (5′ to 3′) using gene fragments (Twist Biosciences) between BsmBI sites with NEBuilder HiFi DNA assembly. Our best performing MutSβ- and MutSα-specific reporters, designs #T5 and #4, respectively (Figures 2D-2G), were cloned into two backbones, pALD-lentiEGFP-A and FUGW (Addgene #14883; generating pYJ461 and pYJ464).

Our MutSα-specific reporters are selectively responsive to loss of MutSα-driven MMR; however, these reporters generally demonstrate lower dynamic range than our MutSβ-specific reporters (Figures 2F, 2G, 3A-3C, S4A, S4B, and S4I). This difference may be caused by higher baseline editing efficiencies in MMR-proficient cell lines with the MutSα-specific reporters, which would impose a more limited ceiling on the maximal fold change that could be observed from perturbing MMR. To improve the dynamic range of our MutSα-specific reporters, future efforts may identify MutSα-specific edits with lower baseline efficiencies or remove evopreQ1, which is known to enhance editing efficiencies, from those reporters.

### Generation of K562 cells with MMR gene disruptions and stable expression of PEmax

K562 PEmax parental cells^65^ and the *MLH1* knockout derivative^56^ (herein, K562 PEmax MLH1-KO) were previously described. We made six additional MMR knockout cell lines in K562 cells for this study: K562 PEmax MSH3-KO (*MSH3* disrupted), K562 PEmax MSH6-KO (*MSH6* disrupted), K562 PEmax MSH2-KO (*MSH2* disrupted), K562 PEmax PMS2-KO (*PMS2* disrupted), K562 PEmax PMS1-KO (*PMS1* disrupted), and K562 PEmax MLH3-KO (*MLH3* disrupted). To generate each of these cell lines, we electroporated K562 PEmax parental cells (∼5.5e5) with 62 pmol of Alt-R S.p. Cas9 Nuclease V3 (Integrated DNA Technologies, 1081058) and 100 pmol of Alt-R CRISPR-Cas9 sgRNA (pre-complexed for 20 min at room temperature) using SE Cell Line 96-well Nucleofector Kit (Lonza, V4SC-1096) with program FF-120, according to manufacturer’s instructions. Sequences of sgRNAs targeted to *MSH3* (Hs.Cas9.MSH3.1.AA), *MSH6* (Hs.Cas9.MSH6.1.AF), *MSH2* (Hs.Cas9.MSH2.1.AA), *PMS2* (Hs.Cas9.PMS2.1.AB), *PMS1*(Hs.Cas9.PMS1.1.AA), and *MLH3* (Hs.Cas9.MLH3.1.AA) are included in Table S2. Seven days later, single cells were isolated by fluorescence-activated cell sorting (BD, FACSAria Fusion). Cells were expanded and those with high percentage of EGFP expression (i.e., similar to the K562 PEmax parental cells) were characterized by amplicon sequencing and western blot analysis (Figures S1D-S1N).

### Self-targeting prime editing screens

Prime editing screens performed with our lYJ003 and lYJ004 self-targeting libraries were conducted in replicates. For each replicate, we transduced 100e6 cells, K562 PEmax parental or MMR knockout derivative cell lines at an MOI of 0.3 using lentivirus prepared as described above (*Lentivirus packaging and transduction*). Prior to transduction, cells were isolated by centrifugation and resuspended in 25 mL complete medium supplemented with 8 µg/mL polybrene (final density of 4e6 cells/mL). Transductions were performed with centrifugation in 6-well plates (16e6 cells/well, 1000 xg for 2 hours at room temperature). After centrifugation, cells were recovered in an 37 °C incubator for 30 mins, and then resuspended to 5e5 cells/mL using complete RPMI medium to a final volume of 200 mL. Cells were cultured in a Multitron (Infors HT) at 37 °C, 85% humidity, 63 rpm, and with 5% (v/v) CO_2_. After 48 hours, 100 mL complete RPMI medium supplemented with 3 µg/mL puromycin (Goldbio, P-600-500) and 0.5% DMSO (1/200 v/v) was added to each culture. Cells were cultured for another 5 days, splitting as needed, and collected for analysis 7 days after transduction. After collection, cells were washed with PBS and stored at -80 °C until sample preparation.

To prepare screen samples for sequencing, cell pellets of ∼100e6 cells were thawed (one pellet per replicate sample). Genomic DNA (gDNA) was then extracted using NucleoSpin Blood XL kit (Macherey-Nagel, 740950.50) according to manufacturer’s instructions, and library elements were amplified for sequencing. We assembled 60-100 PCR reactions per replicate sample (100 µL total per reaction) using NEBNext Ultra II Q5 Master Mix (New England Biolabs, M0544X). Each PCR reaction contained 10 µg gDNA and 0.1 nmol of each of two primers: one complementary to part of the mU6 promoter (oSL059, 5′-AATGATACGGCGACCACCGAGATCTACACGCACAAAAGGAAACTCACCCT-3′) and one complementary to an i7 annealing region on the construct and bearing a standard Illumina i7 index (oYJ023-oYJ034, exact sequences included in Table S4, 5′-CAAGCAGAAGACGGCATACGAGATNNNNNNNNGTGACTGGAGTTCAGACGTGTGCTCTTC-3′). PCR reactions were performed at 98 °C for 1 min, then 22 cycles of [98 °C for 10 s, 65 °C for 30 s, and 72 °C for 45 s], followed by 72 °C for 5 min. PCR products were purified using SPRIselect reagent, quantified by a Qubit 4 Fluorometer, pooled with equal molarity, and evaluated using an Agilent 2100 Bioanalyzer (Agilent, G2939B). Final samples were sequenced using a NovaSeq 6000 S1 300 cycles kit (Illumina, 20028317) with a pair-end read structure of R1 = 184 nt (using oSL055 as a custom sequencing primer), I1 = 8 nt, and R2 = 144 nt. R1 sequenced the pegRNAs, I1 sequenced i7 indexes, and R2 sequenced the target sites and barcodes.

### Generation of RPE-1 hTERT cells with MMR gene disruptions

We made three MMR knockout cell lines in RPE-1 hTERT cells for this study: RPE-1 hTERT MSH3-KO (*MSH3* disrupted), RPE-1 hTERT MSH6-KO (*MSH6* disrupted), RPE-1 hTERT PEmax MSH2-KO (*MSH2* disrupted). To generate each of these cell lines, we electroporated RPE-1 hTERT parental cells (∼2e5) with 76 pmol of Alt-R S.p. Cas9 Nuclease V3 (Integrated DNA Technologies, 1081058) and 125 pmol of Alt-R CRISPR-Cas9 sgRNA (pre-complexed for 20 min at room temperature) using SE Cell Line 96-well Nucleofector Kit (Lonza, V4SC-1096) with program EA-104, according to manufacturer’s instructions. sgRNAs sequences are included in Table S2. Twelve days later, single cells were isolated by fluorescence-activated cell sorting (BD, FACSAria Fusion). Cells were expanded and characterized by amplicon sequencing and western blot analysis (Figures S3C and S3D).

### Ectopic expression of MMR proteins in K562 PEmax cells, related to **Figures 3B**, 3C, and S4C-S4H

We designed rescue constructs that allowed us to express MMR proteins at qualitatively different levels. Our “strong construct” drove MMR gene expression from a UBC promoter. Our “weak construct” drove MMR gene expression from an IRES. For relevant experiments, K562 PEmax parental and MMR knockout cells were transduced with these constructs, each harboring a control gene (emiRFP/TagBFP) or an MMR gene, at an MOI < 0.4, using lentivirus prepared as described above (*Lentivirus packaging and transduction*). Transductions were performed in triplicate for each group. After centrifugation, cells were cultured for 48 hours. Cells were then selected by 3 µg/mL puromycin and expanded daily and, after a week, transduced with MutSα/β-specific reporters and assayed for reporter signals (*MutSα/β-specific reporter assays performed with MMR knockout cells*).

### Western blot analysis

To prepare samples, we isolated ∼5e5 cells by centrifugation and lysed them in 50 μL 1X Bolt^TM^ LDS Sample Buffer (Invitrogen, B0007) and 50 mM Bolt^TM^ Sample Reducing Agent (Invitrogen, B0009). Lysates were incubated at 70 °C for 10 min and then at 85 °C for 3 min with agitation (1250 rpm). To degrade DNA and RNA, lysates were supplemented with 5 mM MgCl_2_ and 1.25 U μL^−1^ GENIUS™ Nuclease (Santa Cruz Biotech, sc-202391) and incubated at 37 °C for 30-45 min with agitation (1250 rpm). Samples were stored at -20 °C prior to further analysis. For SDS-PAGE, samples were thawed, mixed, loaded onto Bolt^TM^ Bis-Tris Plus Mini 4-12% 15-well gels (Invitrogen, NW04125BOX), and run for 40 min at 150 V using MES SDS running buffer (Invitrogen, B0002) with either PageRuler^TM^ Plus Prestained Protein Ladder (Thermo Scientific, 26619) or Precision Plus Protein Dual Color standards (Bio-Rad, 161-0374). Proteins were transferred onto nitrocellulose (Invitrogen, IB23001) using an iBlot^TM^ 2 Dry Blotting System (Invitrogen, IB21001) following manufacturer’s instructions. Resulting membranes were blocked in 5% non-fat dry milk in TBST (Santa Cruz Biotech, sc-2324) for 1 h at room temperature. Proteins were detected using antibodies included in Table S3 using standard procedures.

### Cell electroporation, related to **Figures 2A-2C**, S3B, and S3E

We electroporated plasmids into K562 PEmax cells using the SE Cell Line 96-well Nucleofector Kit (Lonza, V4SC-1096) and program FF-120, following manufacturer’s instructions (Figures 2A and 2B). Typically, ∼5e5 cells were used per reaction with 250 ng pegRNA-expression plasmid and/or 500 ng expression plasmid for MMR proteins or control proteins. Afterward, cells were allowed to recover for 10 min at 37 °C. Cells were then mixed with pre-warmed, antibiotic-free RPMI 1640 medium supplemented with 10% FBS and cultured for 3 days prior to collection. We electroporated plasmids into HAP1 cells using the SE Cell Line 96-well Nucleofector Kit (Lonza, V4SC-1096) and program DZ-113, following manufacturer’s instructions (Figures 2C and S3B). Typically, ∼3e5 cells were used per reaction with 150 ng pegRNA-expression plasmid, 450 ng prime editor-expression plasmid, and/or 300 ng expression plasmid for MMR proteins or control proteins.. Afterward, cells were allowed to recover for 10 min at 37 °C. Cells were then mixed with pre-warmed, antibiotic-free IMDM medium supplemented with 10% FBS and cultured for 3 days prior to collection. We electroporated plasmids into RPE-1 hTERT cells using the SE Cell Line 96-well Nucleofector Kit (Lonza, V4SC-1096) and program EA-104, following manufacturer’s instructions (Figure S3E). In this case, ∼1.5e5 cells were nucleofected with 500 ng prime editor-expression plasmid and 200 ng pegRNA-expression plasmid. Afterward, cells were allowed to recover for 10 min at 37 °C. Cells were then mixed with pre-warmed, antibiotic-free DMEM/F12 medium supplemented with 10% FBS and cultured for 3 days prior to collection. We note that rescue effects in Figures 2B and 2C were incomplete, especially in MSH6-KO cells, likely because MSH6 is highly abundant in many cell types,^103,104^ and ectopic expression may not have reached endogenous expression level. For all experiments listed here, three independent nucleofections were carried out for each experimental group (biological replicates n = 3).

### Targeted amplicon sequencing, related to **Figures 2A-2C**, S1D-S1F, S1J-S1L, S3C, and S4B

We performed amplicon sequencing to characterize K562 and RPE-1 hTERT knockout cell lines and quantify prime editing efficiencies. We first prepared cell lysates by plating ∼150 μL cells into individual wells of 96-well poly-D-lysine coated plates (Greiner Bio-One, 655946, for K562) or regular 96-well plates (Greiner Bio-One, 655180, for RPE-1 hTERT). Cells were incubated overnight (>12 h) to allow plate attachment, and then culture medium was replaced with 1X lysis buffer (∼40-50 μL / well) composed of 10 mM Tris pH 8.0 (Gibco, AM9855G), 0.05% SDS (Invitrogen, 15553027), 25 μg / mL proteinase K (Invitrogen, AM2546), and nuclease-free water (Invitrogen, AM9939). Cells were lysed at 37 °C for 90 min, and resulting lysates were transferred into a PCR plate (Thermo Scientific, AB2400), incubated at 80 °C for 45 min to heat-inactivate proteins, and used for PCR amplification immediately or after storage at -20 °C. Lysates for quantifying prime editing efficiencies were prepared with the following differences: (1) we isolated edited K562 cell populations by centrifugation (200 x g for 5 min at room temperature) and lysed cells with 1X lysis buffer (∼60 μL / 500,000 cells), or (2) we removed medium from adherent cells and lysed directly with 1X lysis buffer (∼40 μL).

Amplicon sequencing libraries were prepared using two rounds of PCR. In the first round (PCR1), targeted loci were amplified with primers containing an annealing region (Tm = 56-64 °C) and an adaptor sequence (5′-ACACTCTTTCCCTACACGACGCTCTTCCGATCT-3′ or 5′-TGGAGTTCAGACGTGTGCTCTTCCGATCT -3′; Table S4). PCR1 reaction mixtures were assembled with 0.1 nmol of each primer, 1.5 µL of cell lysate, 10 µL of NEBNext Ultra II Q5 Master Mix (New England Biolabs, M0544X), and water up to 20 µL. PCR1 reactions were performed at 98 °C for 30 s, then 29-30 cycles of [98 °C for 10 s, 61 °C for 20 s, and 72 °C for 40 s], followed by 72 °C for 2 min. PCR1 products were confirmed by agarose gel electrophoresis. In the second round (PCR2), we extended the adapter region on each amplicon to add P5, i5, P7, and i7 sequences. PCR2 reaction mixtures were assembled with 0.1 nmol of each primer, 1.5 µL of PCR1 product, 10 µL Phusion U Multiplex PCR Master Mix (Thermo Scientific, F564S), and water up to 20 µL. PCR2 reactions were performed at 98 °C for 2 min, then 8 cycles of [98 °C for 10 s, 61 °C for 20 s, and 72 °C for 40 s], followed by 72 °C for 2 min. After amplification, PCR2 products were pooled by volume and purified by gel extraction using Zymoclean Gel DNA Recovery Kit (Zymo Research, D4001). Final products were then quantified by a Qubit 4 Fluorometer (Thermo Scientific, Q33238), examined by Agilent 2100 Bioanalyzer (Agilent, G2939B), and sequenced using a MiSeq Micro (Illumina, MS-103-1002) or Nano (Illumina, MS-103-1001) kit with R1 = 300 nt, i7 = 8 nt, and i5 = 8 nt read structure.

### MutSα/β-specific reporter assays performed with MMR knockout cells, related to **Figures 2F**, 2G, 3A-3C, S4A, S4B, S4I, and S4J

Reporter plasmids were packed into lentivirus and delivered into the cells by lentiviral transduction (*Lentivirus packaging and transduction*). For reporter assays in K562 PEmax cell lines, cells were transduced with the reporter lentivirus at an MOI of ∼1.5 (Day 0). After 2-4 days of culture, cells were mixed and analyzed by flow cytometry (Attune NxT) to determine the live cell number and mCherry positive cell number, which indicates successful reporter transduction. Another aliquot of cells (100 μL) were plated into 96-well white assay plates (Corning, 3917), mixed with equal volume (100 μL) of reconstituted Nano-Glo Luciferase Assay reagents (Promega, N1150), and luminescence signals were read out using a plate reader (Agilent BioTek Synergy HTX, S1LFA-SN or BioTek Cytation 1) using the following parameters: excitation: plug; emission: hole; optics position: top; gain: between 80-120; integration time: 0:00.10 MM:SS.ss; read height: 4.5 mm (for HTX) or 7 mm (for Cytation 1); read speed: normal; delay: 100 msec. For experiments related to Figures 2F, 2G, and S4B, cells were assayed 72 hours post transduction. For experiments related to Figures 3A and S4I, cells were assayed 96 hours post transduction. For experiments related to Figures 3B and 3C, cells were assayed 48 hours post transduction. For experiments related to Figure S4A, cells were assayed at either 72 (x-axis data) or 96 (y-axis data) hours post-transduction. We note that data presented in Figures 3A, S4A, and S4B are from the same experiments.

For reporter assays in RPE-1 hTERT cells (Figure S4J), RPE-1 hTERT parental and derivative MMR knockout cells were transduced with the reporter lentivirus at an MOI of ∼1.5 and expanded for a week. Cells (∼2e5) were then electroporated with 500 ng pCMV-PEmax using the SE Cell Line 96-well Nucleofector Kit (Lonza, V4SC-1096) with program EA-104, according to manufacturer’s instructions. Cells were trypsinized and resuspended in DMEM/F12 medium 2 days after electroporation, and then resuspended cells were analyzed by flow cytometry as above for K562 cells. Another aliquot of cells (100 μL) were plated and assayed also as described above. For these experiments, we normalized the reporter signals to the number of mCherry positive cells per sample before calculating the fold change.

Replicates for these experiments were established as follows: for experiments related to Figures 2F and 2G, a single transduction was performed for each experimental group and cells were sampled post-transduction and assayed in independent wells as replicates (n = 2 or 3). For experiments related to Figures 3A-3C, S4A, S4B, S4I, and S4J, three independent transductions were performed (n = 3) and each was independently assayed.

### MutSβ-specific reporter assays for siRNA or cDNA overexpression, related to Figures S4K and S4L

MutSβ-specific reporter constructs were transduced into the K562 PEmax cells as described above (*MutSα/β-specific reporter assays performed with MMR knockout cells*). Cells (5e5) were nucleofected with indicated amounts of siRNA (Figure S4K; Table S2) or 250 ng indicated plasmids (Figure S4L) 2 days after transduction using the SE Cell Line 96-well Nucleofector Kit (Lonza, V4SC-1096) with program FF-120 according to manufacturer’s instructions. After 3 days of recovery, cells were analyzed by flow cytometry and NanoGlo luciferase reagents, as described above. For these experiments, we normalized the resulting luminescence to numbers of mCherry positive cells in each sample, as described above.

### MutSα/β-specific reporter assays performed with drug treatments in 96-well plates, related to Figures 3D, 3E, 4B, 4C, S5A, S5C-S5G, S5K, S5L, S6A-S6E

For reporter assays in K562 PEmax cells, cells were transduced with reporter constructs at an MOI of ∼1.5 (Figures 3D, 3E, 4B, 4C, S5A, S5E, S5G, S5K, S5L, and S6A-S6C) or at indicated MOIs (Figure S5F). Transduced cells were treated with DMSO (control) or indicated small molecules at the indicated concentrations 2 days post transduction, cultured for another 2 days (Figures 3D right, 3E, 4B, 4C, S5A, S5E-S5G, S5K, S5L, and S6A-S6C) or for indicated amounts of time (Figure 3D left) and then analyzed with NanoGlo luciferase reagents. For the experiments depicted in Figure S5G left, different luciferase reagents were also used (Renilla-Glo for Renilla Luc and ONE-Glo for Firefly Luc, Promega, E2710 and E6110 respectively).

For reporter assays in HAP1, HeLa, U-2 OS, Lenti-X 293T, and HCT 116 cells (Figures S5C and S5D), cells were transduced with reporter constructs at an MOI of 1-2. Transduced cells were then cultured and split as needed for 4-5 days, and then transfected with pCMV-PEmax-EGFP or pCMV-PE7-EGFP, by TurboFectin 8.0 (OriGene, TF81001) for HAP1, TransIT-HeLaMONSTER (mirus bio, MIR2904) for HeLa, Lipofectamine 2000 (Invitrogen, 11668027) for U-2 OS and Lenti-X 293T, and Lipofectamine LTX (Invitrogen, 15338030) for HCT 116, all following manufacturers’ protocols. Transfected cells were treated with DMSO (control) or cmpd1 2 days later, plated into 96-well white assay plates with clear bottom (Corning, 3610), cultured for another 2 days, and then analyzed with NanoGlo luciferase reagents also as described above.

For reporter assays in RPE-1 hTERT cells (Figures S6D and S6E), cells were transduced with PE7 at an MOI of ∼0.3. Cells were expanded after transduction and miRFP680 positive cells (with prime editor constructs) were sorted by fluorescence-activated cell sorting (BD, FACSAria Fusion). Cells were expanded again and then transduced with the MutSβ-specific reporter lentiviruses at an MOI of ∼1.5. Transduced cells were treated with small molecules at indicated concentrations 2 days post transduction, cultured in 96-well white assay plates with clear bottoms (Corning, 3610) for another 2 days, and then analyzed by NanoGlo luciferase reagent, also as described above.

Replicates for these experiments were established as follows: Cells were mixed with indicated concentrations of drugs and split into separate wells of 96-well plates. Luminescence was examined separately for each well and resulting data were treated as individual replicates. Experiments in Figures 4B, 4C, S5A, S5K, S5L, and S6A-S6C were performed with 3 such replicates; experiments in Figures 3D, 3E, S5C-S5F, S5G left, S6D, and S6E were performed with 4 such replicates; and experiments Figure S5G (right, 96-well plates) were performed with 18 such replicates. We note that, for small molecule-related experiments, we did not normalize luminescence to numbers of mCherry positive cells in each sample.

### Cell viability assays (standard throughput), related to Figure S5B

Cells were treated with drugs and 2 days later were transferred into wells of 96-well white assay plates (Corning, 3917), mixed with equal volume (100 μL) of CellTiter-Glo® 2.0 Cell Viability Assay reagents (Promega G9241), and the luminescence signals were read out by a plate reader (Agilent BioTek Synergy HTX, S1LFA-SN or BioTek Cytation 1) as described above (*MutSα/β-specific reporter assays performed with MMR knockout cells*). Reported cell viability measurements (i.e., fold changes of live cells of drug treated versus DMSO controls) were calculated by normalizing resulting luminescence signals to the mean of luminescence from DMSO-treated cells.

### Composition of small molecule libraries

Small molecule libraries were provided by the Princeton University Small Molecule Screening Center. Altogether, we screened two libraries: the bioactive collection, which consisted of 5,923 small molecules, and the general collection, which consisted of 47,780 individual small molecules and 4,480 pooled mixture with 4 stereoisomers in each well. The total number of small molecules screened was therefore 5,923 + 47,780 + 4,480*4 = 71,623 small molecules.

### Primary small molecule screen, related to **Figure**s 3F, 3G, and S5H-S5J

#### Preparing small molecules

To aliquot compounds for screening, small molecules from each collection were added individually to wells of white, flat-bottom 384-well plates (Thermo Scientific, 164610) at either 0.02 μL or 0.04 μL per well (from 10 mM stock solutions dissolved in DMSO) by an Echo acoustic liquid handler (Beckman Coulter, Labcyte Echo 550). On each plate, we left the first and last two columns (1, 2, 23, 24) empty and, immediately before screening, added DMSO or cmpd1 to those wells (DMSO, 1 and 23; cmpd1, 2 and 24; 0.02 μL per well). Bioactive collection small molecules were screened at a final concentration of 10 and 20 μM, while general collection compounds were screened at a final concentration of 10 μM. As an internal standard, we ran a control plate with only titrated cmpd1 alongside each batch of plates in the screen.

#### Preparing and processing cells

K562 PEmax cells were transduced with the MutSβ-specific reporter at an MOI of ∼1.5. After 48 hours of recovery, transduced cells were analyzed by flow cytometry (AttuneNxT, Invitrogen, A24858) and transduction at the targeted MOI was confirmed by mCherry expression. Cells were then diluted to 5e5 cells/mL in fresh RPMI complete medium and dispensed into compound-containing 384-well plates at 20 μL/well using a Multidrop Combi reagent dispenser (Thermo Scientific, 5840300) with small tube metal tip dispensing cassette (Thermo Scientific, 24073295). Cells were allowed to grow for 48 hours and were then assayed for luminescence using Nano-Glo Luciferase Assay reagents (Promega, N1150) dispensed at 20 μL per well. Plates rested at room temperature for 5-10 min and then luminescence was read out using a plate reader (Agilent BioTek Synergy HTX, S1LFA-SN) with the following parameters: excitation, plug; emission, hole; optics position, top; gain, 120; integration time, 0:00.20 MM:SS.ss; read height, 1.00 mm. Resulting data were exported into .xlsx files using the Agilent Gen5 software for processing and analysis (QUANTIFICATION AND STATISTICAL ANALYSIS). We note that, for the primary small molecule screen, only one replicate was performed.

### Secondary screening of small molecules and individual testing of analogs and resynthesized hits in high-throughput

Hits from the bioactive collection, resynthesized small molecules, and compound analogs were examined in high-throughput with procedures similar to those used in primary screening, with the following exceptions: (1) compounds in these tests were evaluated across a range of concentrations and with biological replicates, and (2) compounds in these tests were examined with both MutSα- and MutSβ-specific reporters. Hits from the bioactive collection (Table S8) were dispensed in volumes of 2.5 nL, 5 nL, 10 nL, 20 nL, 40 nL, and 80 nL into individual wells of white, flat-bottom 384-well plates (2 wells per condition) from 10 mM stock solutions dissolved in DMSO to reach final concentrations of 1.25 μM to 40 μM after subsequent addition of 20 μL of cells. Other small molecules tested in 384-well plates (Figures 4A, 4D-4F, and S6F-S6K) were dispensed in volumes of 2.5 nL, 5 nL, 10 nL, 15 nL, 20 nL, and 40 nL into individual wells of white, flat-bottom 384-well plates (3 wells per condition) from 10 mM stock solutions dissolved in DMSO to reach final concentrations of 1.25 μM to 20 μM after subsequent addition of 20 μL of cells. When testing small molecules in this format, cells without reporters were often similarly processed and used to examine compound toxicities. Cell viability measurements were made by adding 20 μL of CellTiter-Glo 2.0 Cell Viability Assay reagent (Promega, G9242) to the cells and quantifying luminescence on a plate reader (Agilent BioTek Synergy HTX, S1LFA-SN) with the following parameters: excitation, plug; emission, hole; optics position, top; gain, 120; integration time, 0:00.20 MM:SS.ss; read height, 1.00 mm. Other procedures were the same as described above (*Primary small molecule screen*).

### Reverse transcription-quantitative polymerase chain reaction (RT-qPCR)

For RT-qPCR experiments, total RNA was extracted using Monarch® Total RNA Miniprep Kit (NEB, T2010S) and quantified by NanoDrop. Total RNA was then used for RT-qPCR reactions using Power SYBR™ Green RNA-to-CT™ 1-Step Kit (Applied Biosystems, 4389986), following manufacturer’s instructions. Each 20 μL reaction contained 50 ng of RNA, 8 pmol of each of the forward and reverse primers (Table S4), 0.16 μL of 125X RT enzyme mix, and 10 μL of Power SYBR Green Master Mix. Reactions were then run on a ViiA 7 Real-Time PCR System (Applied Biosystem) with the following program: 48 °C for 30 min, 95 °C for 10 min, then 40 cycles of [95 °C for 15 s, 65 °C for 1 min], followed by 95 °C for 15 s, 60 °C for 1 min, increase to 95 °C at 0.05 °C/s, and 95 °C for 15 s. ΔΔCt values^105^ were calculated and converted to percentage of expression level. For these experiments, biological replicates represent three independent samples of compound-treated cells, and for each of those replicates, 4 technical replicates using the same reaction master mix were aliquoted into 4 wells in the assay plate were also included.

### DNA intercalation assay, related to Figure S7C

SPR measurements were performed on a Cytiva Biacore 8K+ (Danaher Life Sciences LLC, USA) using Series S Streptavidin-coated biochips (Danaher Life Sciences LLC, USA). Measurements were performed in the following assay buffer: 50 mM HEPES, pH 7.5, 150 mM KCl, 5 mM MgCl_2_, 0.5 mM TCEP, 500 μM ATP, 0.05% (v/v) Tween-20, 0.05% BSA, and 1% DMSO. After standard chip conditioning according to the manufacturer’s instructions, 1.5 μM 5′-biotinylated 38 bp (CAG)_2_ DNA was captured in the assay buffer to a target level of 2500 RU. For this experiment, the sample and chip compartment were set to T = 20 °C. All compounds were diluted in 100% DMSO using a Biomek FXP (Beckman Coulter Inc, USA) from 10 mM stocks in a 3-fold dilution series to 1.52 µM. The compound dilution series was transferred to 384-micro well plates as 1:100 dilutions in the assay buffer using an Integra Viaflo 384 system (INTEGRA Biosciences GmbH, Germany). Test compounds were then injected into the SPR flowcell. Contact duration of association was 120 s, dissociation duration 120 s at a flow rate of 30 µL/min with a data collection rate of 10 Hz. To calculate the binding signals, a 5 seconds window, 115-120 s after initial injection of compounds, was taken to calculate an average signal (background signal from DNA binding to the flow cell subtracted) for each small molecule at each concentration.

### SPR assay, related to **Figures** 5A, 5B, S7D, and S7E

SPR measurements were performed with equipment described above (*DNA intercalation assay*). Measurements were performed in the following assay buffer: 50 mM HEPES, pH 7.5, 150 mM KCl, 5 mM MgCl_2_, 0.5 mM TCEP, 100 μM ATP, 0.05% (v/v) Tween-20. After standard chip conditioning according to the manufacturer’s instructions, 0.35 nM 5’-biotinylated 38 bp (CAG)_2_ DNA was captured in running buffer without DMSO to a target level of 25 RU for MutSβ measurements; analogously for MutSα measurements, 5′-biotinylated 38 bp G-T mismatch DNA was captured. For these experiments, the sample and chip compartment were set to T = 20°C. All compounds were diluted in 100% DMSO using a Biomek FXP (Beckman Coulter Inc, USA) from 10 mM stocks in a 3-fold dilution series to 4.57 µM. The compound dilution series were transferred to 384-micro well plates as 1:100 dilutions in assay buffer with either 15 nM MutSβ or 50 nM MutSα using an Integra Viaflo 384 system (INTEGRA Biosciences GmbH, Germany). After 30 minutes of incubation, MutSα or MutSβ only control and mixtures with the compound dilution series were injected according to the SPR schema. Contact duration of association was 120 s (0-120 s, schematized in Figure S7D), dissociation duration 240 s at a flow rate of 30 µL/min with a data collection rate of 10 Hz. After dissociation, surface regeneration was performed by injection of 0.1% SDS for 30 seconds. To calculate the binding signals, the “analyte binding late” region of the data (5 seconds window, 115-120 s, schematized in Figure S7E) was taken to calculate an average signal (background signal from DNA binding to the flow cell subtracted) for each sample at each concentration. The calculated binding signals were then normalized to the results from no compound control (0 µM).

### FRET assay, related to **Figures** 5C, 5D, and S7F

Assay was performed with or without ATP as described^81^ with some modifications: The assay buffer was modified to contain 1 mM TCEP instead of DTT, and 100 nM protein and 100 µM ATP was used. The assay was carried out in a 384-well low volume round bottom black plate (Corning 4514) in 8 µL total assay volume. Assay materials were prepared as follows: First, a compound serial dilution plate was generated via Biomek FX Laboratory Automation Workstation with 12-point, 3-fold dilution starting from 20 mM top concentration of each compound in DMSO in a 384-well plate (Greiner). Second, MutSβ was added at 4 µL to the assay plate with matrix pipette (Thermo scientific). Third, compounds were dipped at 58 nL from the compound serial dilution plate to the assay plate with CyBio pintool such that the maximum concentration of compounds on the assay plate was 145 µM. For MutSβ+DNA or free DNA controls without compound, an equivalent amount of DMSO was dipped instead of compounds from the compound plate. Samples were then mixed briefly and incubated for 30 min at room temperature. Meanwhile probe plate containing Cy3-Cy5 labeled 70 bp (CAG)_1_ loop DNA or such DNA plus ATP was prepared and 4 µL of probe (labeled DNA +/- ATP) was transferred via Integra Viaflo 384 from the probe plate to the assay plate such that the final labeled DNA concentration in the assay was 5 nM. Samples were mixed and fluorescence of labeled DNA was recorded in donor (Ex 545 nm/Em 590 nm) and acceptor channels (Ex 545 nm/Em 680 nm) in kinetic mode over 70 min. The FRET ratio (fluorescence in acceptor channel / fluorescence in donor channel) was recorded for steady state.

### Plasmid-based mismatch repair assay, related to **Figures 5E**, 5F, and S7G

Preparations of the proteins and plasmid DNA substrates were described previously.^69^ Reactions were carried out in a final volume of 15 µL in buffer containing 25 mM Tris-acetate (pH 7.5), 0.1 mg/mL recombinant albumin (New England Biolabs, B9200S), 5 mM MgCl₂, and 2 mM ATP. Each reaction contained 80 ng of the 3.2 kb plasmid DNA substrate pSam (T)₄. No additional NaCl or KCl was added. The only salt present came from the recombinant protein storage buffer, giving a final NaCl concentration of approximately 20 mM. Protein storage buffer was added as needed to ensure the same final NaCl concentration in all samples. Where indicated, inhibitors were used at 100 µM. MutSβ (75 nM) or FAN1 (75 nM) was added on ice, and reactions were then incubated with the indicated inhibitor for 5 min at 37 °C. Next, 75 nM MutLγ or MutLα, 75 nM RFC, and 75 nM PCNA (where indicated) were added, and the reactions were incubated for a further 45 min at 37 °C. Reactions were stopped by adding 5 µL STOP solution (150 mM EDTA, 2% SDS, 30% glycerol, and 0.01% bromophenol blue) together with 1 µL Proteinase K (Roche, 18 mg/mL), followed by incubation at 50 °C for 1 h. Products were separated on 1% agarose gels containing GelRed (Biotium) in TAE buffer (40 mM Tris, 20 mM acetate, and 1 mM EDTA) for 90 min at 6 V/cm. Gels were imaged using a Quantum gel imager (Vilber). Band intensities were quantified with ImageJ2 (NIH) and data were plotted and analyzed in Prism 10 (GraphPad).

### Cell-based CAG repeat expansion assay, related to Figures 5G-5I

RPE-1 hTERT (CAG)_129_ cells carry a stably integrated mHTT gene exon 1 (originally introduced with lentiviral transduction). We plated these cells at 2.5e5 cells per well in 6 well plates and added the indicated compounds at indicated concentrations. After 48 hours, initial samples (Day 0) were collected. Cells were then subcultured for 21 additional days, by splitting 1:4 every 3 to 4 days. Compounds were replenished during reseeding. Samples were taken every 7 days. Genomic DNA was then extracted from each sample using the DNeasy Blood & Tissue kit (Qiagen, 69509), and CAG repeat tracts within the integrated mHTT gene exon 1 were amplified by PCR using 100 ng of genomic DNA, AmpliTaq Gold 360 Master Mix (Applied Biosystems, 4398876), a 6-FAM labelled forward primer (5′-CCTTCGAGTCCCTCAAGTCCTT-3′), and a reverse primer (5′-CGGCTGAGGCAGCAGCGGCTGT-3′) with the following program: 95 °C for 10 min, then 35 cycles of [95 °C for 30 s, 58 °C for 30 s, and 72 °C for 90 s], followed by 72 °C for 7 min. PCR products were denatured in Hi-Di™ Formamide (Thermo Scientific, 4440753) and analyzed by capillary electrophoresis using the 3730xl DNA Analyzer with MapMarker® 1000 labelled with ROX (Eurogentec, MW-0195-80ROX). Raw data from capillary electrophoresis was analysed using the R package “trace: Tandem Repeat Analysis by Capillary Electrophoresis” (https://cran.r-project.org/web/packages/trace/index.html). Briefly, ladder standard peak was aligned with CAG repeat tract peaks and modal repeat length was called by determining the modal peak above 20% of the maximum peak signal. Per day changes in modal CAG repeat lengths were calculated as follows: (modal CAG repeats length on Day 21 - modal CAG repeats length on Day 0) / 21.

### Cell cycle analysis by Click EdU assay, related to Figure S7H

RPE-1 hTERT (CAG)_129_ cells were treated with compounds as described above (*Cell-based CAG repeat expansion assay*) and collected for cell cycle analysis on Day 14 (drug treatment for 16 days total). Cells were incubated with EdU for 1 hour at 37 °C. Cells were then fixed and EdU was labeled using the Click-iT™ EdU Alexa Fluor™ 647 Flow Cytometry Assay Kit (Thermo Scientific, C10419) following manufacturer’s instructions. DNA was labeled by incubating resulting EdU-labeled cells in 2 µg/mL propidium iodide and 100 µg/mL RNase A. For analysis, 20,000 events were recorded per sample by flow cytometry (Cytoflex, Beckman technology).

## QUANTIFICATION AND STATISTICAL ANALYSIS

### Analysis of targeted amplicon sequencing

To quantify prime editing outcomes from targeted amplicon sequencing, we used a previously reported approach.^65^ Briefly, we aligned sequencing reads to corresponding reference sequences (Table S4) with CRISPResso2^106^ (2.2.11) using HDR batch mode. We defined the intended outcome as the expected HDR amplicon sequence (“--expected_hdr_amplicon_seq”) and used the following settings: the minimum average read quality was 30 (“--min_average_read_quality 30”), and the quantification window size was 20 (“--quantification_window_size 20”) and centered at the pegRNA nick (“--quantification_window_center −3”). We discarded reads with indels (i.e., unwanted editing outcomes, “--discard_indel_reads”). We defined intended editing efficiencies as the fractions of reads containing intended edits without indels and calculated those efficiencies as follows: (number of non-discarded HDR-aligned reads) / (number of reads that aligned all amplicons) X 100%.

### Analysis of self-targeting prime editing libraries (lYJ003 and lYJ004)

Plasmid libraries were cloned and sequenced as described above (*Cloning of HEK3 (lYJ003) and FANCF (lYJ004) self-targeting libraries*). Reads from validation sequencing were expected to contain the following:

Read 1 (5′ to 3′): spacer (20 nt), sgRNA scaffold (76 nt), extension (variable for each edit), and polyT (5′-TTTTTTTCC-3′) sequences.

Read 2 (5′ to 3′): target site (90 nt), barcode (12 nt) flanked by XhoI and BamHI recognition sequences.

Quality assessment was performed on read pairs with average Phred ≥ 30 in two multipart steps. <u>Step 1:</u> <u>Assigning reads to library elements</u>. (1) Spacer and scaffold regions (20 + 76 nt) were searched on read 1 to match the expected sequences, regardless of the starting position. Reads with no spacer-scaffold region found were discarded. (2) The polyT sequence was searched downstream of the spacer-scaffold region on read 1, and we extracted the sequence in between the spacer-scaffold and polyT. Reads for which the extracted sequence perfectly matched a designed extension sequence were retained and assigned to a library element while others were discarded. (3) For reads assigned to a library element, the corresponding barcode was extracted from between the XhoI and BamHI recognition sequences on read 2. Extracted barcodes were compared to expected sequences, and reads for which the extracted barcode matched the expected barcode up to one Levenshtein distance were retained. <u>Step 2: Verification of target site</u> <u>sequences</u>. (1) After confidently assigning reads to library elements, we verified the target site sequences (90 nt) by extracting them from the beginning of read 2. Reads were retained if the extracted target site sequence differed from the expected target site sequence by fewer than 4 bp.

To assess the uniformity of these libraries, we calculated a Gini coefficient using read counts for each element, after removing any recombined elements.

### Analysis of self-targeting prime editing screens (performed with lYJ003 and lYJ004)

For analysis of screen data, reads were assigned to library elements essentially as described for Step 1 in *Analysis of self-targeting prime editing libraries (lYJ003 and lYJ004*). For this analysis, a separate FASTQ file was created for each library element containing read 2 sequences retained after Step 1. These files were processed using Cutadapt^107^ to trim off the restriction enzyme recognition sequences and barcode, leaving the 90 nt target sequence. Reads shorter than 10 bp after trimming were discarded. Trimmed reads were then analyzed using CRISPResso2 (version 2.3.1) with the following parameters:

--exclude_bp_from_left 0 --exclude_bp_from_right 0 --amplicon_seq [reference/unedited sequence] -g [protospacer sequence] -e [HDR/edited sequence] -q 30 -w 0 --discard_indel_reads.

This analysis used trimmed read 2 sequences for alignment and classification into three categories: reference, HDR, or ambiguous. A minimum average Phred quality score of 30 was required, and reads falling below that score were removed entirely from any downstream calculation. The window size was set to 0 so that each read would be classified based on its full sequence. Reads containing indels were not considered HDR events but were retained in the total read count (defined below as the number of reads that aligned to all amplicons). Note that in this analysis, we use “indel(s)” to refer to unintended insertions and/or deletions introduced during prime editing, library preparation, or sequencing, which is different from intended insertion or deletion prime edits.

As enough sequencing depth is required for accurate calculation of editing efficiencies, we discarded all elements with less than 500 reads aligning to all amplicons.

CRISPResso2 generated a file for each library element with two rows containing read counts aligned to the (1) reference sequence and (2) the HDR sequence (which in this case, is the reference sequence containing the intended prime edit). Columns of interest were Reads_aligned_all_amplicons (total number of reads that are able to align to either the reference sequence or HDR sequence), Modified (number of reads aligned to the reference or HDR sequence with mismatches), Unmodified (number of reads aligned to the reference or HDR sequence without mismatches), and Discarded (number of reads aligned to the reference or HDR sequence that contain indels, with or without mismatches). Editing efficiencies were calculated as: (number of unmodified HDR-aligned reads) / (number of reads that aligned all amplicons).

During analysis, we considered that library elements may obtain edits from unassigned pegRNAs when multiple constructs enter the same cell, and moreover, recombination events and sequencing errors may produce similar effects. We therefore estimated a “background editing rate” for each library element that was designed to make an edit (i.e., “editing elements”) using elements that were designed not to introduce any edit (i.e., “negative control elements”). That is, we determined the frequencies at which modifications from the former set of elements unexpectedly appeared across the latter. To calculate these frequencies, we first applied CRISPResso2 across our negative controls. We performed this analysis without the -e parameter, as no HDR sequences were expected for those elements. Similar to editing elements, negative controls with fewer than 500 reads after library assignment were discarded.

For each negative control element, CRISPResso2 generated a table summarizing all read patterns observed among assigned read 2 sequences, recorded the number of read counts per pattern, and indicated read patterns containing an indel (denoted as DISCARDED_Reference). We then further assigned these read patterns into categories using custom scripts as follows: (1) <u>NaN_mutation</u>: a read pattern containing substitutions that does not perfectly match any editing element in the library; (2) <u>NaN_indel</u>: a read pattern containing indels that does not perfectly match any editing element in the library; (3) <u>Unedited</u>: a read pattern that perfectly matches the reference sequence; (4) <u>Assigned_others</u>: a read pattern that perfectly matches the HDR sequence corresponding to an editing element in the library.

We used the Assigned_others category to calculate the frequencies at which modifications from each editing element appeared across all negative controls, and defined that as the background editing rate for the corresponding element. As a stringent filter, we excluded any library element whose intended editing efficiency, in any replicate of screen data from any of the MMR knockout cell lines, was < 3 standard deviations above the mean of its background editing rate. This filter removed 28 elements from downstream analysis for HEK3 Screens 1 and 2 and removed 8 elements for the FANCF Screen.

### Analysis of the small molecule screen

Screen data were compressed from raw read .xlsx files into a summary .xlsx file for statistical analysis. For each plate a Z′-factor was calculated based on the means (μ) and standard deviations (σ) of raw luminescence signals from negative (DMSO) and positive (cmpd1) control wells:

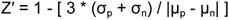

where p and n denote positive and negative controls, respectively.

These Z′-factors were used to assess and maintain data quality throughout our screen. That is, plates with a Z′ < 0.2 (rounded to one decimal place) were rerun until a Z′ >= 0.2 was obtained. Out of 204 plates in our screen, 9 (4.41%) had an initial Z′ < 0.2, and the first rerun of all 9 substantially improved the Z′ to satisfy the Z′ >= 0.2 requirement. For clarity, only plates with a Z′ >= 0.2 were used for downstream analysis. For technical reasons, positive controls were not added to the first plate in the screen (index number 1), and we therefore excluded that plate from downstream analysis.

We next used a relative reporter activation metric to score compounds in our screen, herein called “fraction of reporter activation relative to cmpd1 control”. This metric was calculated from raw luminescence signals on a per-plate basis as:

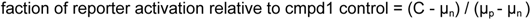

where C represents the raw luminescence of the test compound and p and n denote positive and negative controls, respectively.

This metric directly compares the activities of each test compound to signals from our positive and negative controls, such that a score of 0 would mean that cells treated with test compound produced signals equivalent to those treated with vehicle (DMSO), and a score of 1 would mean that cells treated with test compound produced signals equivalent to those treated with cmpd1. We also calculated this metric for each well of DMSO-treated cells and determined hits as compounds that scored > 3 standard deviations from the mean of all such wells, across all plates in the screen. Compounds scoring above this threshold were then reevaluated in secondary testing using resynthesized compounds or new stocks from vendors (*Secondary screening of small molecules and individual testing of resynthesized hits and analogs in high-throughput*).

### Analysis from secondary testing of small molecules in high-throughput

For determining MutSβ-specific reporter effects in these experiments, we used procedures essentially the same as described above (*Analysis of the small molecule screen*). In this analysis, a small molecule significantly activated the MutSβ-specific reporter if the corresponding fraction of reporter activation relative to cmpd1 control was ≥3 standard deviations from the mean of DMSO controls (0.075) at either 10 or 20 µM (Table S8). For determining MutSα-specific reporter effects, we used an alternative metric, as we lacked a positive control for calculating a relative metric. Instead, we calculated a Z-score:

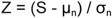

where S represents the raw luminescence of the test compound and n denotes negative control.

In this analysis, a small molecule significantly activated the MutSα-specific reporter if the corresponding Z-score was >=3 (Table S8).

