## Supplementary Figures for "Prime editing enables high-throughput identification and characterization of selective MutSβ inhibitors"

#### SUPPLEMENTARY FIGURES AND LEGENDS

### FIGURE S1

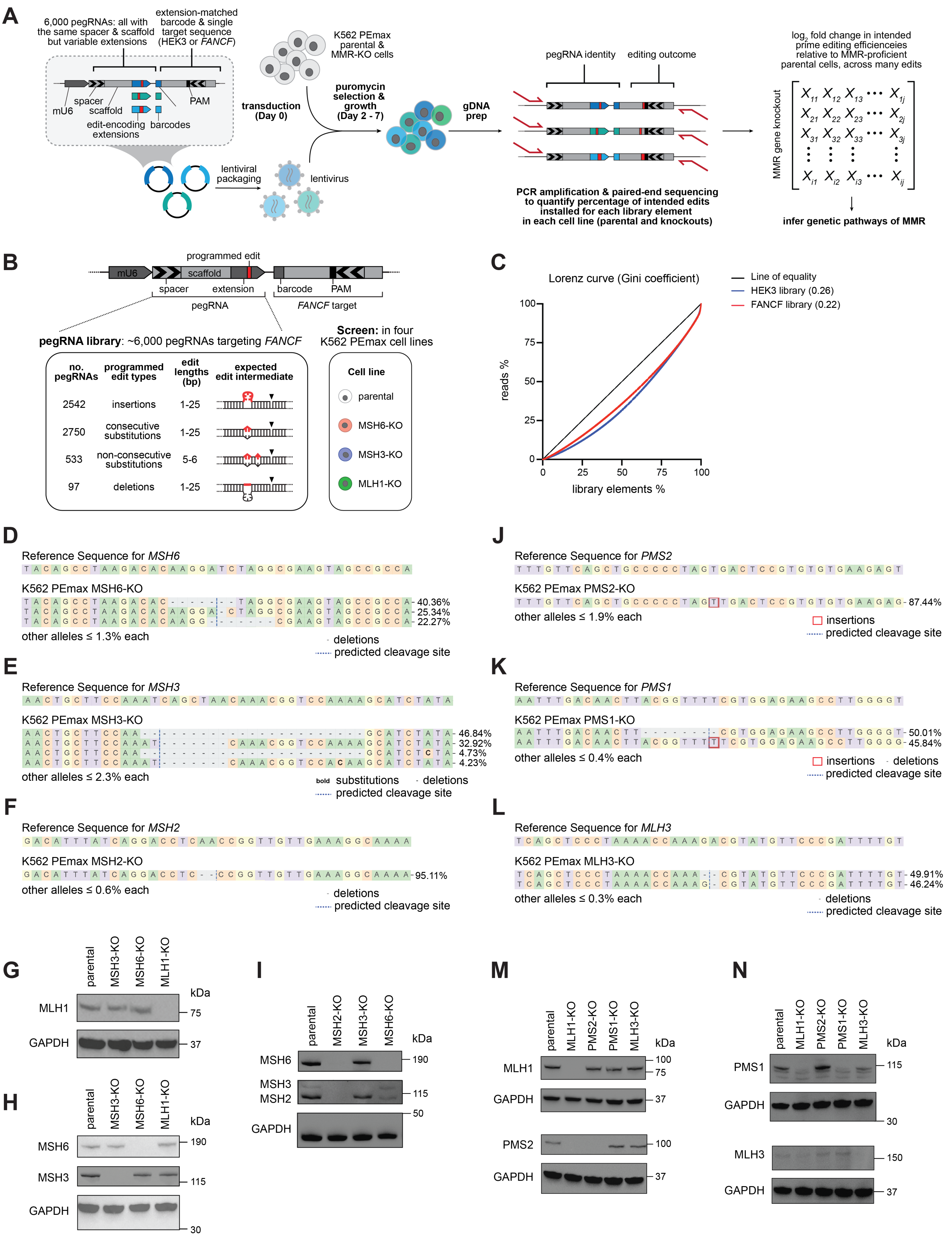
A
6,000 pegRNAs: all with the same spacer & scaffold but variable extensions
extension-matched barcode & single target sequence (HEK3 or FANCF)
spacer
mU6
scaffold
PAM
edit-encoding extensions
barcodes
lentiviral packaging
lentivirus
transduction (Day 0)
K562 PEmax parental & MMR-KO cells
puromycin selection & growth (Day 2 - 7)
gDNA prep
pegRNA identity
editing outcome
PCR amplification & paired-end sequencing to quantify percentage of intended edits installed for each library element in each cell line (parental and knockouts)
log2 fold change in intended prime editing efficienceies relative to MMR-proficient parental cells, across many edits
MMR gene knockout
infer genetic pathways of MMR
B
programmed edit
mU6
scaffold
spacer
extension
pegRNA
barcode
PAM
FANCF target
pegRNA library: ~6,000 pegRNAs targeting FANCF
no. pegRNAs
programmed edit types
edit lengths (bp)
expected edit intermediate
2542 insertions 1-25
2750 consecutive substitutions 1-25
533 non-consecutive substitutions 5-6
97 deletions 1-25
Screen: in four K562 PEmax cell lines
Cell line
parental
MSH6-KO
MSH3-KO
MLH1-KO
C
Lorenz curve (Gini coefficient)
Line of equality
HEK3 library (0.26)
FANCF library (0.22)
reads %
library elements %
D
Reference Sequence for MSH6
K562 PEmax MSH6-KO
40.36%
25.34%
22.27%
other alleles ≤ 1.3% each
deletions
predicted cleavage site
E
Reference Sequence for MSH3
K562 PEmax MSH3-KO
46.84%
32.92%
4.73%
4.23%
other alleles ≤ 2.3% each
bold substitutions
deletions
predicted cleavage site
F
Reference Sequence for MSH2
K562 PEmax MSH2-KO
95.11%
other alleles ≤ 0.6% each
deletions
predicted cleavage site
J
Reference Sequence for PMS2
K562 PEmax PMS2-KO
87.44%
other alleles ≤ 1.9% each
insertions
predicted cleavage site
K
Reference Sequence for PMS1
K562 PEmax PMS1-KO
50.01%
45.84%
other alleles ≤ 0.4% each
insertions
deletions
predicted cleavage site
L
Reference Sequence for MLH3
K562 PEmax MLH3-KO
49.91%
46.24%
other alleles ≤ 0.3% each
deletions
predicted cleavage site
G
parental
MSH3-KO
MSH6-KO
MLH1-KO
kDa
MLH1
75
GAPDH
37
H
parental
MSH3-KO
MSH6-KO
MLH1-KO
kDa
MSH6
190
MSH3
115
GAPDH
30
I
parental
MSH2-KO
MSH3-KO
MSH6-KO
kDa
MSH6
190
MSH3
MSH2
115
50
GAPDH
M
parental
MLH1-KO
PMS2-KO
PMS1-KO
MLH3-KO
kDa
MLH1
100
75
GAPDH
37
PMS2
100
GAPDH
37
N
parental
MLH1-KO
PMS2-KO
PMS1-KO
MLH3-KO
kDa
PMS1
115
GAPDH
30
MLH3
150
GAPDH
37

**Figure S1. Design of prime editing screens and characterization of MMR knockout cell lines, related to Figure 1**

(A) Schematic of prime editing screens. Library constructs were packaged into lentivirus and delivered into K562 PEmax parental and derivative MMR knockout cell lines. After 5 days of puromycin selection (Day 2 to 7), genomic DNA (gDNA) was extracted and the integrated constructs were PCR amplified and sequenced to quantify the editing outcomes, which were then used to infer MMR genetic pathways.

(B) Schematic depicting prime editing library using the *FANCF* target site and related screen. Self-targeting lentiviral constructs (top) were used for delivery of the target sequence and pegRNAs programming many DNA mispairs to that site (bottom left). Screens were performed in K562 PEmax parental cells and derivative cell lines with core MMR genes knocked out (bottom right). Arrows within the depicted construct indicate directionality of the pegRNA spacer and matched protospacer within the target.

(C) Lorenz curves from sequencing the HEK3 and FANCF plasmid libraries. Library elements are defined as pegRNA-target site pairs. Gini coefficients for each library included in parentheses.

(D-F) Characterization of indels at targeted loci in indicated K562 PEmax knockout cell lines: MSH6-KO cells in D, MSH3-KO cells in E, and MSH2-KO cells in F. Percentages represent the fraction of reads with the indicated allele observed in amplicon sequencing. Reference Sequence is from parental cells. Analysis performed with CRISPResso2.

(G-I) Western blot analyses of K562 PEmax cells and derivative knockout clones: immunoblotting of MLH1 in G, MSH6 and MSH3 in H, and MSH6, MSH3, and MSH2 in I. Groups of images are from the same membrane, each of which was simultaneously incubated with indicated antibodies, except for H and I. Membrane in H was coblotted for MSH6 and GAPDH prior to stripping and reblotting for MSH3. Membrane in I was coblotted for MSH2 and GAPDH prior to stripping and reblotting for MSH6 and then for MSH3.

(J-L) Characterization of indels at targeted loci in indicated K562 PEmax knockout cell lines: PMS2-KO cells in J, PMS1-KO cells in K, MLH3-KO cells in L. Percentages represent the fraction of reads with the indicated allele observed in amplicon sequencing. Reference Sequence is from parental cells. Analysis performed with CRISPResso2. We note that characterization of the MLH1-KO cell line was previously reported.[56]

(M-N) Western blot analyses of K562 PEmax parental cells and derivative knockout clones: immunoblotting of MLH1 and PMS2 in M, and PMS1 and MLH3 in H. Groups of images are from the same membrane, each of which was simultaneously blotted with the indicated antibodies.

### FIGURE S2

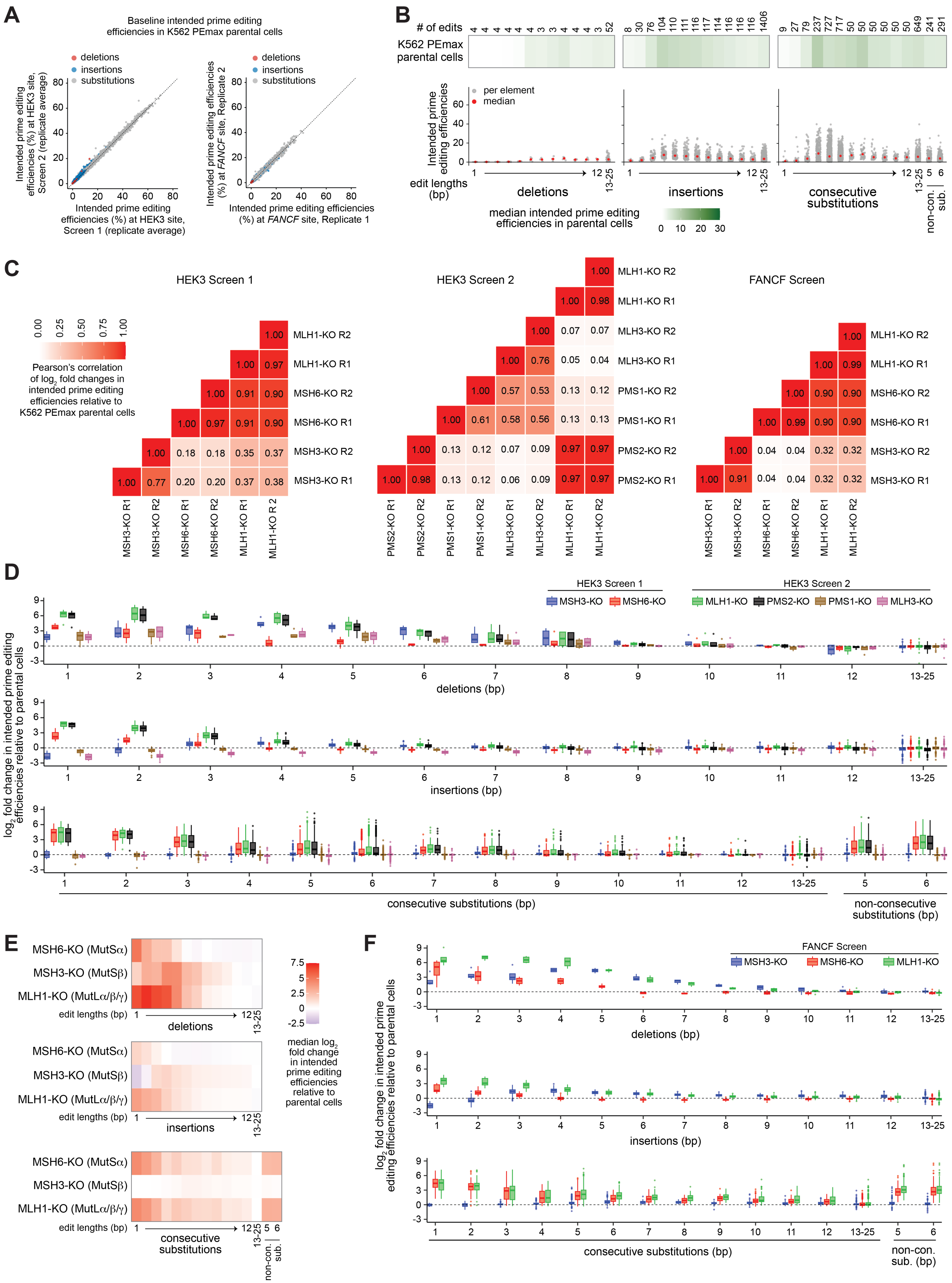
A
Baseline intended prime editing efficiencies in K562 PEmax parental cells
deletions
insertions
substitutions
Intended prime editing efficiencies (%) at HEK3 site, Screen 2 (replicate average)
Intended prime editing efficiencies (%) at HEK3 site, Screen 1 (replicate average)
Intended prime editing efficiencies (%) at FANCF site, Replicate 2
Intended prime editing efficiencies (%) at FANCF site, Replicate 1
B
### of edits
4 4 4 4 4 4 3 3 4 4 4 3 52
8 30 76 104 110 111 116 117 114 116 116 116 1406
9 27 79 237 727 717 50 50 50 50 50 649 241 291
K562 PEmax parental cells
per element
median
Intended prime editing efficiencies
edit lengths (bp)
deletions
insertions
consecutive substitutions
non-con. sub.
13-25
median intended prime editing efficiencies in parental cells
0 10 20 30
C
HEK3 Screen 1
HEK3 Screen 2
FANCF Screen
Pearson's correlation of log2 fold changes in intended prime editing efficiencies relative to K562 PEmax parental cells
0.00 0.25 0.50 0.75 1.00
MLH1-KO R2: 1.00
MLH1-KO R1: 1.00 0.97
MSH6-KO R2: 1.00 0.91 0.90
MSH6-KO R1: 1.00 0.97 0.91 0.90
MSH3-KO R2: 1.00 0.18 0.18 0.35 0.37
MSH3-KO R1: 1.00 0.77 0.20 0.20 0.37 0.38
MSH3-KO R1, MSH3-KO R2, MSH6-KO R1, MSH6-KO R2, MLH1-KO R1, MLH1-KO R 2
MLH1-KO R2: 1.00
MLH1-KO R1: 1.00 0.98
MLH3-KO R2: 1.00 0.07 0.07
MLH3-KO R1: 1.00 0.76 0.05 0.04
PMS1-KO R2: 1.00 0.57 0.53 0.13 0.12
PMS1-KO R1: 1.00 0.61 0.58 0.56 0.13 0.13
PMS2-KO R2: 1.00 0.13 0.12 0.07 0.09 0.97 0.97
PMS2-KO R1: 1.00 0.98 0.13 0.12 0.06 0.09 0.97 0.97
PMS2-KO R1, PMS2-KO R2, PMS1-KO R1, PMS1-KO R2, MLH3-KO R1, MLH3-KO R2, MLH1-KO R1, MLH1-KO R2
MLH1-KO R2: 1.00
MLH1-KO R1: 1.00 0.99
MSH6-KO R2: 1.00 0.90 0.90
MSH6-KO R1: 1.00 0.99 0.90 0.90
MSH3-KO R2: 1.00 0.04 0.04 0.32 0.32
MSH3-KO R1: 1.00 0.91 0.04 0.04 0.32 0.32
MSH3-KO R1, MSH3-KO R2, MSH6-KO R1, MSH6-KO R2, MLH1-KO R1, MLH1-KO R2
D
HEK3 Screen 1
HEK3 Screen 2
MSH3-KO MSH6-KO MLH1-KO PMS2-KO PMS1-KO MLH3-KO
log2 fold change in intended prime editing efficiencies relative to parental cells
deletions (bp)
insertions (bp)
consecutive substitutions (bp)
non-consecutive substitutions (bp)
E
MSH6-KO (MutSα)
MSH3-KO (MutSβ)
MLH1-KO (MutLα/β/γ)
edit lengths (bp)
deletions
insertions
consecutive substitutions
non-con. sub.
7.5 5 2.5 0 -2.5
median log2 fold change in intended prime editing efficiencies relative to parental cells
F
FANCF Screen
MSH3-KO MSH6-KO MLH1-KO
log2 fold change in intended prime editing efficiencies relative to parental cells
deletions (bp)
insertions (bp)
consecutive substitutions (bp)
non-con. sub. (bp)

**Figure S2. Additional analysis of prime editing screens, related to Figure 1**

(A) Comparison of intended prime editing efficiencies at the HEK3 (left) or *FANCF* (right) target site in K562 PEmax parental cells between two different screens (HEK3 Screens 1 and 2) or between two replicates of one screen (FANCF Screen). Each dot represents an element in the prime editing library, colored by edit type (red, deletions; blue, insertions; grey, substitutions). Black dashed lines indicate y = x.

(B) Heatmaps (top) show median intended prime editing efficiencies at the *FANCF* target site in K562 PEmax parental cells grouped based on edit length and type, as in Figure 1D. The number of prime edits in each group is included. Dot plots (bottom) show individual (grey dots) and median (red dots) intended prime editing efficiencies within each group.

(C) Comparisons of $\log_2$ fold changes in intended prime editing efficiencies relative to parental cells between different samples from the HEK3 Screen 1 (left), HEK3 Screen 2 (middle), and FANCF Screen (right). Data represent Pearson's correlation coefficients.

(D) Box plots depict $\log_2$ fold changes in intended prime editing efficiencies relative to parental cells from HEK3 Screens 1 and 2, grouped by edit types (top, deletions; middle, insertions; bottom, substitutions) and separated by edit lengths (increasing from left to right) and cell lines (indicated with colors).

(E) Heatmaps show median $\log_2$ fold changes in intended prime editing efficiencies for each knockout cell line at the *FANCF* target site, stratified by prime edit length and type, as in Figure 1D.

(F) Box plots depict $\log_2$ fold changes in intended prime editing efficiencies relative to parental cells from the FANCF Screen, grouped by edit types (top, deletions; middle, insertions; bottom, substitutions) and separated by edit lengths (increasing from left to right) and cell lines (indicated with colors).

### FIGURE S3

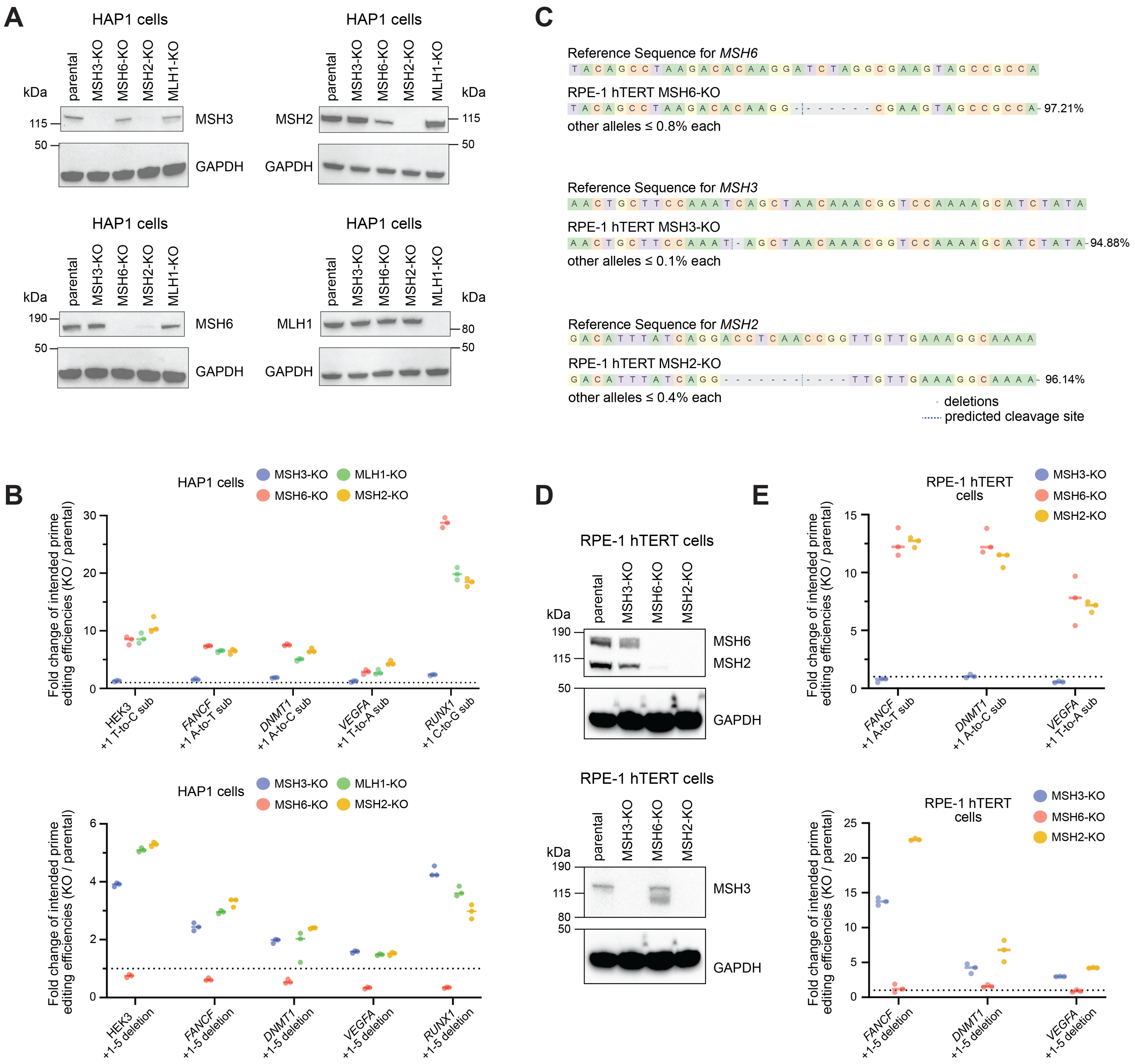
A
HAP1 cells
parental
MSH3-KO
MSH6-KO
MSH2-KO
MLH1-KO
kDa
115
50
MSH3
GAPDH
MSH2
190
MSH6
MLH1
80
C
Reference Sequence for MSH6
TACAGCCTAAGACACAAGGATCTAGGCGAAGTAGCCGCCA
RPE-1 hTERT MSH6-KO
TACAGCCTAAGACACAAGG-------CGAAGTAGCCGCCA- 97.21%
other alleles ≤ 0.8% each
Reference Sequence for MSH3
AACTGCTTCCAAATCAGCTAACAAACGGTCCAAAAGCATCTATA
RPE-1 hTERT MSH3-KO
AACTGCTTCCAAAT-AGCTAACAAACGGTCCAAAAGCATCTATA- 94.88%
other alleles ≤ 0.1% each
Reference Sequence for MSH2
GACATTTATCAGGACCTCAACCGGTTGTTGAAAGGCAAAA
RPE-1 hTERT MSH2-KO
GACATTTATCAGG-----------TTGTTGAAAGGCAAAA- 96.14%
other alleles ≤ 0.4% each
- deletions
predicted cleavage site
B
HAP1 cells
MSH3-KO
MLH1-KO
MSH6-KO
MSH2-KO
Fold change of intended prime editing efficiencies (KO / parental)
HEK3 +1 T-to-C sub
FANCF +1 A-to-T sub
DNMT1 +1 A-to-C sub
VEGFA +1 T-to-A sub
RUNX1 +1 C-to-G sub
HEK3 +1-5 deletion
FANCF +1-5 deletion
DNMT1 +1-5 deletion
VEGFA +1-5 deletion
RUNX1 +1-5 deletion
D
RPE-1 hTERT cells
parental
MSH3-KO
MSH6-KO
MSH2-KO
kDa
190
115
50
80
MSH6
MSH2
MSH3
GAPDH
E
RPE-1 hTERT cells
MSH3-KO
MSH6-KO
MSH2-KO
Fold change of intended prime editing efficiencies (KO / parental)
FANCF +1 A-to-T sub
DNMT1 +1 A-to-C sub
VEGFA +1 T-to-A sub
FANCF +1-5 deletion
DNMT1 +1-5 deletion
VEGFA +1-5 deletion

**Figure S3. Further characterization of MutSα/β resolving prime edits (1 bp substitutions and 5 bp deletions) and additional MMR knockout cell lines, related to Figure 2**

(A) Western blot analyses of HAP1 parental cells and derivative knockout clones: immunoblotting of MSH3 (upper left), MSH2 (upper right), MSH6 (lower left), and MLH1 (lower right). Groups of images are from the same membrane, each of which was simultaneously blotted with the indicated antibodies.

(B) Fold changes in intended prime editing efficiencies for 1 bp substitution edits (top) or 5 bp deletion edits (bottom) at indicated genomic loci in indicated cell lines relative to parental cells. Groups of dots represent data from n = 3 biological replicates. pegRNA and PEmax editor plasmids were delivered by nucleofection.

(C) Characterization of indels at targeted loci in indicated RPE-1 hTERT knockout cell lines: MSH6-KO (top), MSH3-KO (middle), and MSH2-KO (bottom). Percentages represent the fraction of reads with the indicated allele observed in amplicon sequencing. Reference Sequence is from parental cells. Analysis performed with CRISPResso2. We note that reference sequences are identical to those in Figures S1D-S1F, as relevant sequences are the same in RPE-1 hTERT and K562 cells.

(D) Western blot analyses of RPE-1 hTERT parental cells and derivative knockout clones: immunoblotting of MSH6 and MSH2 (upper) and MSH3 (lower). Groups of images from the same membrane, each of which was simultaneously blotted with the indicated antibodies.

(E) Fold changes in intended prime editing efficiencies for 1 bp substitution edits (top) or 5 bp deletion edits (bottom) at indicated genomic loci in indicated cell lines relative to parental cells. Groups of dots represent data from n = 3 biological replicates. pegRNA and PEmax editor plasmids were delivered by nucleofection.

#### FIGURE S4

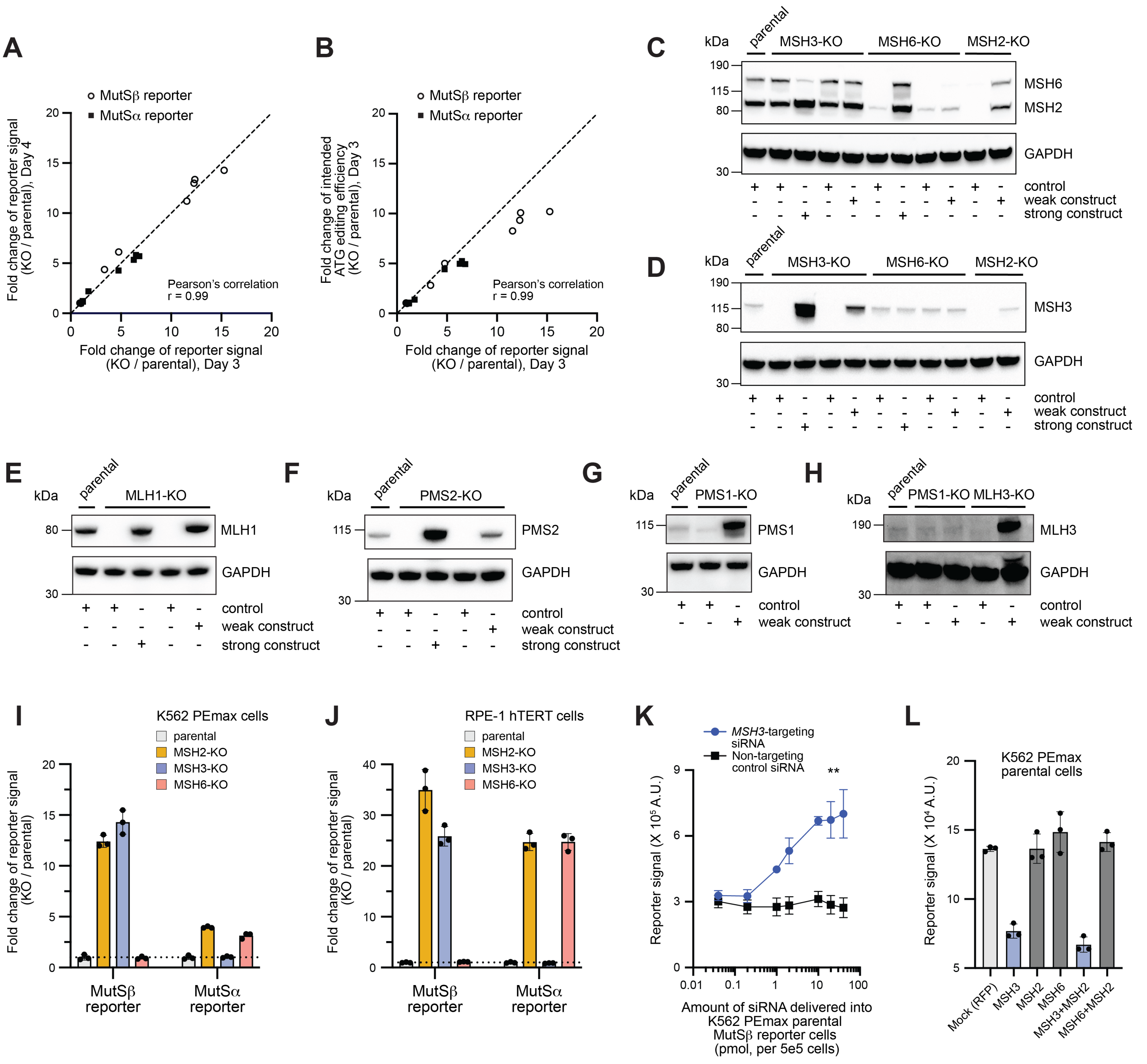

A
MutSβ reporter
MutSα reporter
Fold change of reporter signal (KO / parental), Day 4
Fold change of reporter signal (KO / parental), Day 3
Pearson's correlation r = 0.99
B
Fold change of intended ATG editing efficiency (KO / parental), Day 3
C
kDa
parental
MSH3-KO
MSH6-KO
MSH2-KO
MSH6
MSH2
GAPDH
control
weak construct
strong construct
D
MSH3
E
MLH1-KO
MLH1
F
PMS2-KO
PMS2
G
PMS1-KO
PMS1
H
MLH3-KO
MLH3
I
K562 PEmax cells
parental
MSH2-KO
MSH3-KO
MSH6-KO
Fold change of reporter signal (KO / parental)
MutSβ reporter
MutSα reporter
J
RPE-1 hTERT cells
K
MSH3-targeting siRNA
Non-targeting control siRNA
Reporter signal (X 10^5 A.U.)
Amount of siRNA delivered into K562 PEmax parental MutSβ reporter cells (pmol, per 5e5 cells)
L
K562 PEmax parental cells
Reporter signal (X 10^4 A.U.)
Mock (RFP)
MSH3
MSH2
MSH6
MSH3+MSH2
MSH6+MSH2

**Figure S4. Further genetic validation of MutSα/β-specific reporters, related to Figure 3**

(A) Comparison of MutSα/β-specific reporter activations across K562 PEmax MMR knockout cell lines 3 and 4 days post delivery of each reporter by lentiviral transduction. Dots represent fold changes in luminescence normalized to parental cells presented as means of n = 3 biological replicates. Data from Day 4 (y-axis) were also used in Figure 3A. Black dashed line indicates y = x.

(B) Comparison of MutSα/β-specific reporter activations across K562 PEmax MMR knockout cell lines and fold changes in intended ATG prime editing efficiencies determined by sequencing 3 days post delivery of each reporter by lentiviral transduction. Dots represent fold changes in luminescence normalized to parental cells presented as means of n = 3 biological replicates. MutSα/β-specific reporter activation data (x-axis) were also used in A. Black dashed line indicates y = x.

(C-H) Western blot analyses of K562 PEmax parental cells and derivative knockout clones after transduction of a weak or strong rescue construct matched to the knocked out gene or transduction of a control (Methods): immunoblotting of MSH6 and MSH2 in C, MSH3 in D, MLH1 in E, PMS2 in F, PMS1 in G, and MLH3 in H. Groups of images are from the same membrane, each of which was simultaneously blotted with the indicated antibodies.

(I) Validation of MutSβ- and MutSα-specific reporters (primary designs, #T5 and #4, respectively) using an alternative lentiviral backbone in K562 PEmax parental cells and indicated knockout cell lines. Dots represent fold changes in luminescence, bars represent means of n = 3 biological replicates, and error bars indicate standard deviations.

(J) Validation of our primary MutSβ- and MutSα-specific reporters in RPE-1 hTERT parental cells and indicated knockout cell lines. Dots represent fold changes in luminescence, bars represent means of n = 3 biological replicates, and error bars indicate standard deviations.

(K) Validation of our primary MutSβ-specific reporter in K562 PEmax parental cells after nucleofection with an *MSH3*-targeting or non-targeting siRNA. Dots represent raw luminescence signals presented as means of n = 3 biological replicates and error bars indicate standard deviations. Two-way ANOVA was used for statistical analysis (**, $p < 0.01$).

(L) MutSβ-specific reporter activation in K562 PEmax parental cells after nucleofection with cDNA expression plasmids carrying the indicated genes. Dots represent raw luminescence signals, bars represent means of n = 3 biological replicates, and error bars indicating standard deviations.

### FIGURE S5

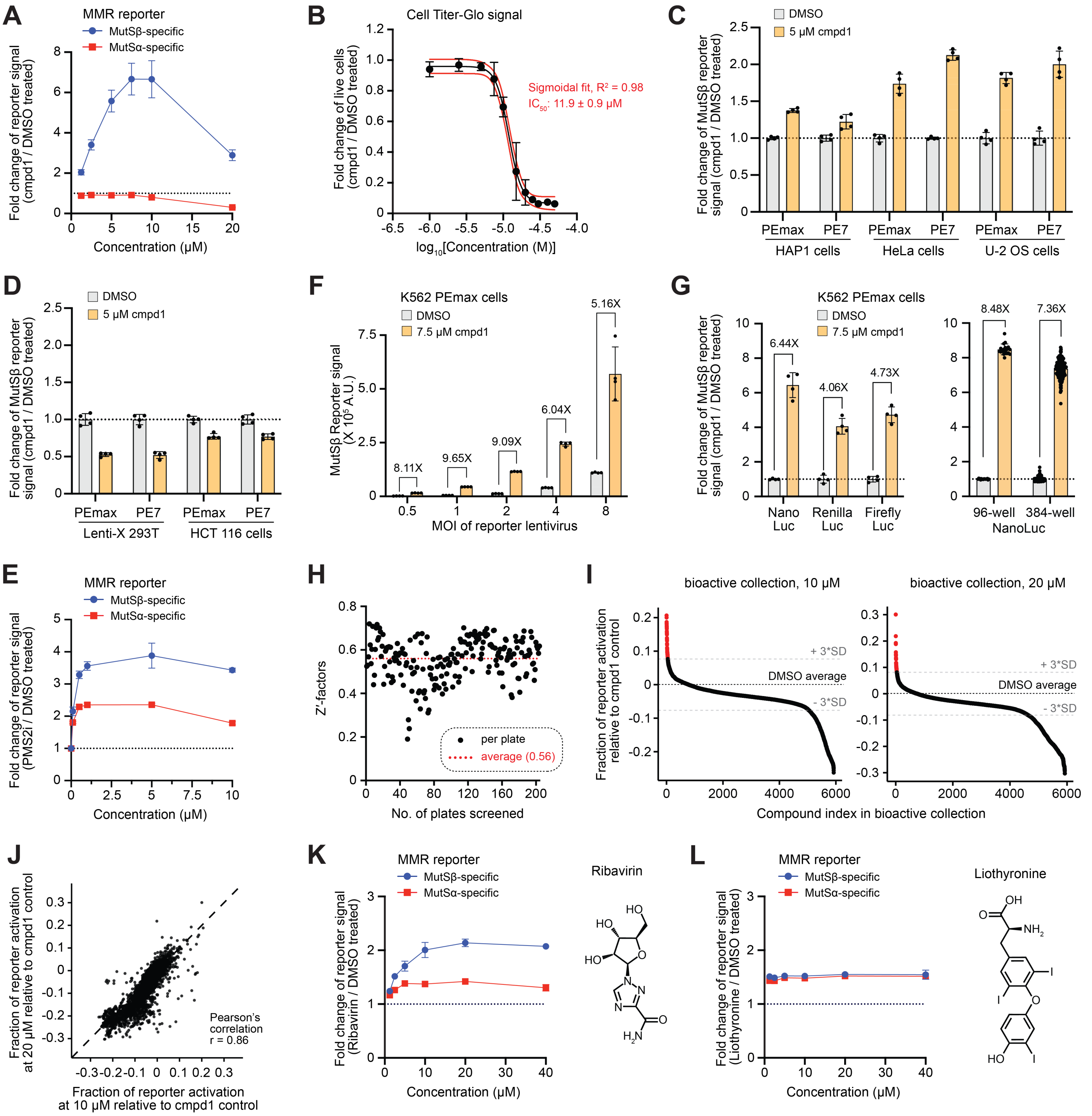

A
MMR reporter
MutSβ-specific
MutSα-specific
Fold change of reporter signal (cmpd1 / DMSO treated)
Concentration (μM)
B
Cell Titer-Glo signal
Fold change of live cells (cmpd1 / DMSO treated)
Sigmoidal fit, $R^2$ = 0.98
$IC_{50}$: 11.9 ± 0.9 μM
$log_{10}$[Concentration (M)]
C
DMSO
5 μM cmpd1
Fold change of MutSβ reporter signal (cmpd1 / DMSO treated)
PEmax
PE7
HAP1 cells
HeLa cells
U-2 OS cells
D
DMSO
5 μM cmpd1
Fold change of MutSβ reporter signal (cmpd1 / DMSO treated)
PEmax
PE7
Lenti-X 293T
HCT 116 cells
F
K562 PEmax cells
DMSO
7.5 μM cmpd1
MutSβ Reporter signal (X $10^5$ A.U.)
8.11X
9.65X
9.09X
6.04X
5.16X
MOI of reporter lentivirus
G
K562 PEmax cells
DMSO
7.5 μM cmpd1
Fold change of MutSβ reporter signal (cmpd1 / DMSO treated)
6.44X
4.06X
4.73X
8.48X
7.36X
Nano Luc
Renilla Luc
Firefly Luc
96-well
384-well
NanoLuc
E
MMR reporter
MutSβ-specific
MutSα-specific
Fold change of reporter signal (PMS2i / DMSO treated)
Concentration (μM)
H
Z'-factors
per plate
average (0.56)
No. of plates screened
I
bioactive collection, 10 μM
bioactive collection, 20 μM
Fraction of reporter activation relative to cmpd1 control
+ 3*SD
DMSO average
- 3*SD
Compound index in bioactive collection
J
Fraction of reporter activation at 20 μM relative to cmpd1 control
Pearson's correlation r = 0.86
Fraction of reporter activation at 10 μM relative to cmpd1 control
K
MMR reporter
MutSβ-specific
MutSα-specific
Fold change of reporter signal (Ribavirin / DMSO treated)
Concentration (μM)
Ribavirin
L
MMR reporter
MutSβ-specific
MutSα-specific
Fold change of reporter signal (Liothyronine / DMSO treated)
Concentration (μM)
Liothyronine

**Figure S5. Establishing a scalable platform for the identification and characterization of MMR inhibitors with our MutSα/β-specific reporters, related to Figure 3**

(A) Plot depicts dose-dependent effects of cmpd1 on MutSβ-specific (blue line) and MutSα-specific (red line) reporter activation in K562 PEmax cells relative to the mean of DMSO controls. Dots represent fold changes in luminescence presented as means of n = 3 biological replicates and error bars indicate standard deviations.

(B) Characterization of cmpd1 toxicity in K562 PEmax cells. Cell Titer-Glo was used to determine the relative number of live cells after treatment with varying concentrations of cmpd1 or treatment with DMSO. Dots represent fold changes in luminescence presented as means of n = 3 biological replicates relative to the mean of DMSO controls and error bars indicate standard deviations. Red curves indicate sigmoidal fit.

(C-D) Effect of cmpd1 on our primary MutSβ-specific reporter in MMR-proficient (C) or MMR-deficient (D) cell lines after 48 hour drug treatments. Dots represent fold changes in luminescence, bars represent means of n = 4 biological replicates, and error bars indicate standard deviations.

(E) Plot depicts dose-dependent effects of PMS2i on MutSβ-specific (blue line) and MutSα-specific (red line) reporter activation in K562 PEmax cells relative to the mean of DMSO controls. Dots represent fold changes in luminescence presented as means of n = 3 biological replicates and error bars indicate standard deviations.

(F) MutSβ-specific reporter activation by 7.5 µM of cmpd1 or DMSO in K562 PEmax cells across a range of MOIs used for reporter transduction. Dots represent raw luminescence signals, bars represent means of n = 4 biological replicates, and error bars indicate standard deviations.

(G) Effect of cmpd1 on MutSβ-specific reporters engineered with NanoLuc, Renilla luciferase (Renilla Luc), or Firefly luciferase (Firefly Luc) indicator genes (left) or on our primary MutSβ-specific reporter in 96- or 384-well plates (right). Experiments performed in K562 PEmax parental cells. Dots represent fold changes in luminescence, bars represent means of n = 4 (left) or 18 (right, 96-well plates) or 192 (right 384-well plates) biological replicates, and error bars indicate standard deviations.

(H) Per-plate Z'-factors from small molecule screening. Dots represent per plate scores and the red line indicates the mean of those scores.

(I) Results from screening the bioactive collection of small molecules at 10 µM (left) or 20 µM (right) using our primary MutSβ-specific reporter (n = 1). Dots represent fractions of reporter activation relative to cpmd1 controls, defined as background-subtracted luminescence from individual wells normalized to the mean of background-subtracted luminescence from on-plate cmpd1 controls, such that 0 represents average results from DMSO controls (black dashed line) and 1 represents average results from cmpd1 controls (Methods). Grey dashed lines indicate 3 standard deviations from the DMSO average (+/- 3*SD). Hits (Table S8) are colored in red.

(J) Comparison of results from screening the bioactive collection of small molecules at 10 µM and 20 µM. Dots represent fractions of reporter activation relative to cpmd1 controls, as described in I.

(K-L) Plots depict dose-dependent effects of Ribavirin (K) or Liothyronine (L) on MutSβ-specific (blue lines) and MutSα-specific (red lines) reporter activation in K562 PEmax cells relative to the mean of DMSO controls. Dots represent fold changes in luminescence presented as means of n = 3 biological replicates and error bars indicate standard deviations. The structure of each small molecule is depicted (insets).

### FIGURE S6

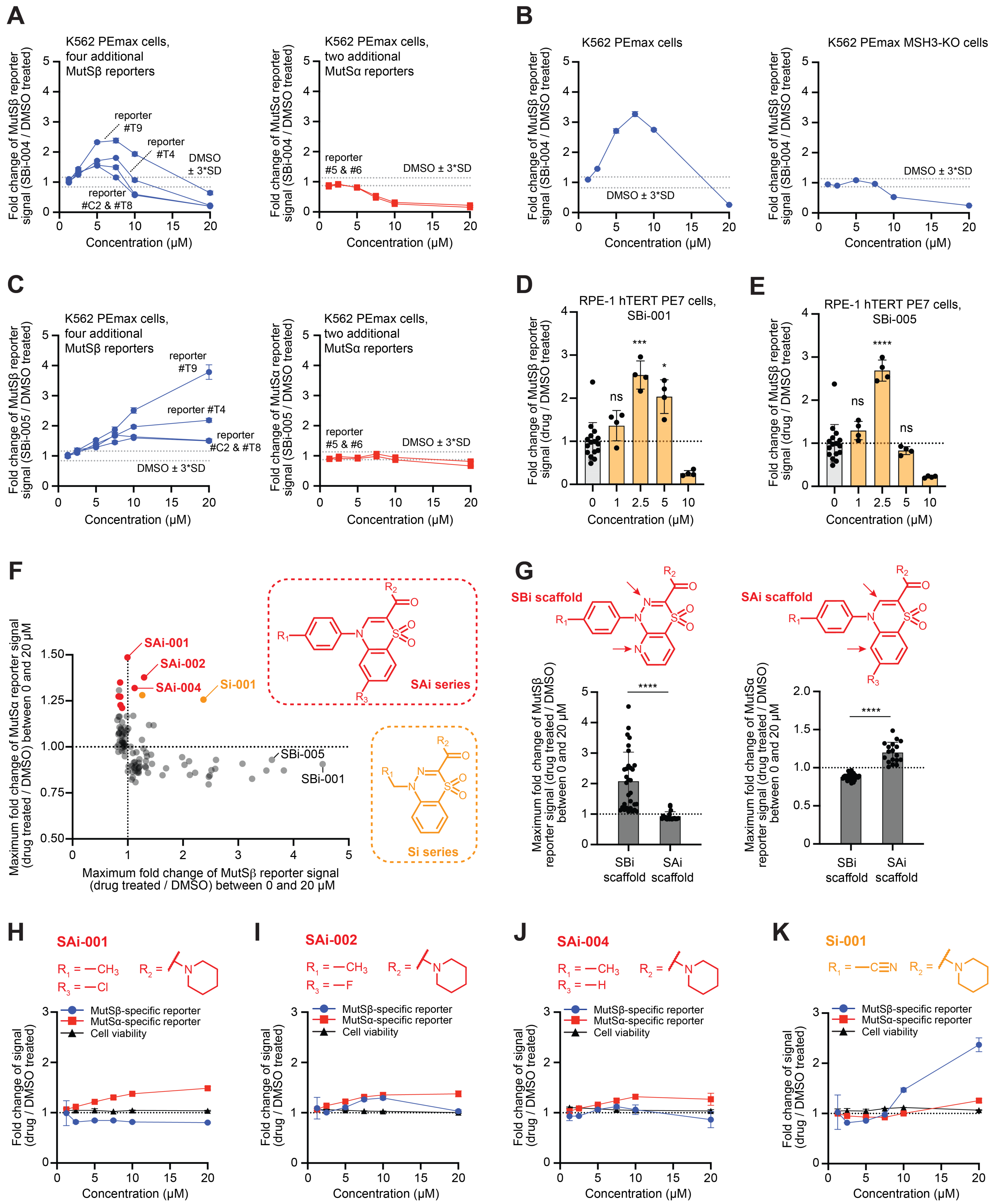
A
K562 PEmax cells, four additional MutSβ reporters
Fold change of MutSβ reporter signal (SBi-004 / DMSO treated)
reporter #T9
reporter #T4
DMSO ± 3*SD
reporter #C2 & #T8
Concentration (μM)
K562 PEmax cells, two additional MutSα reporters
Fold change of MutSα reporter signal (SBi-004 / DMSO treated)
reporter #5 & #6
DMSO ± 3*SD
Concentration (μM)
B
K562 PEmax cells
Fold change of MutSβ reporter signal (SBi-004 / DMSO treated)
DMSO ± 3*SD
Concentration (μM)
K562 PEmax MSH3-KO cells
Fold change of MutSβ reporter signal (SBi-004 / DMSO treated)
DMSO ± 3*SD
Concentration (μM)
C
K562 PEmax cells, four additional MutSβ reporters
Fold change of MutSβ reporter signal (SBi-005 / DMSO treated)
reporter #T9
reporter #T4
reporter #C2 & #T8
DMSO ± 3*SD
Concentration (μM)
K562 PEmax cells, two additional MutSα reporters
Fold change of MutSα reporter signal (SBi-005 / DMSO treated)
reporter #5 & #6
DMSO ± 3*SD
Concentration (μM)
D
RPE-1 hTERT PE7 cells, SBi-001
Fold change of MutSβ reporter signal (drug / DMSO treated)
ns
***
*
Concentration (μM)
E
RPE-1 hTERT PE7 cells, SBi-005
Fold change of MutSβ reporter signal (drug / DMSO treated)
ns
****
ns
Concentration (μM)
F
SAi-001
SAi-002
SAi-004
Si-001
SBi-005
SBi-001
Maximum fold change of MutSα reporter signal (drug treated / DMSO) between 0 and 20 μM
Maximum fold change of MutSβ reporter signal (drug treated / DMSO) between 0 and 20 μM
SAi series
Si series
G
SBi scaffold
SAi scaffold
Maximum fold change of MutSβ reporter signal (drug treated / DMSO) between 0 and 20 μM
****
SBi scaffold
SAi scaffold
Maximum fold change of MutSα reporter signal (drug treated / DMSO) between 0 and 20 μM
****
SBi scaffold
SAi scaffold
H
SAi-001
R1 = —CH3
R3 = —Cl
R2 =
I
SAi-002
R1 = —CH3
R3 = —F
R2 =
J
SAi-004
R1 = —CH3
R3 = —H
R2 =
K
Si-001
R1 = —C≡N
R2 =
MutSβ-specific reporter
MutSα-specific reporter
Cell viability
Fold change of signal (drug / DMSO treated)
Concentration (μM)

**Figure S6. Additional cell-based validation of candidate MutSβ inhibitors and further SAR analysis, related to Figure 4**

(A) Plots depict dose-dependent effects of SBi-004 on activation of additional MutSβ-specific reporters (left, blue lines) and additional MutSα-specific reporters (right, red lines) in K562 PEmax cells relative to the mean of DMSO controls. Dots represent fold changes in luminescence presented as means of n = 3 biological replicates, error bars indicate standard deviations, and dashed lines indicate 3 standard deviations from the mean of DMSO controls.

(B) Plots depict dose-dependent effects of SBi-004 on MutSβ-specific reporter activation in K562 PEmax parental (left) or MSH3-KO (right) cells relative to the mean of DMSO controls. Dots represent fold changes in luminescence presented as means of n = 3 biological replicates, error bars indicate standard deviations, and dashed lines indicate 3 standard deviations from the mean of DMSO controls.

(C) Plots depict dose-dependent effects of SBi-005 on activation of additional MutSβ-specific reporters (left, blue lines) and additional MutSα-specific reporters (right, red lines) in K562 PEmax cells relative to the mean of DMSO controls. Dots represent fold changes in luminescence presented as means of n = 3 biological replicates, error bars indicate standard deviations, and dashed lines indicate 3 standard deviations from the mean of DMSO controls.

(D-E) Effects of SBi-001 (D) and SBi-005 (E) on our primary MutSβ-specific reporter across different concentrations in RPE-1 hTERT cells stably expressing prime editor PE7. Dots represent fold changes in luminescence, bars represent means of n = 4 biological replicates, and error bars indicate standard deviations. Cells with 0 concentration were treated with DMSO. Ordinary one-way ANOVA was used for statistical analysis followed by Dunnett's multiple comparisons test (****, adjusted $p$ value < 0.0001; ***, adjusted $p$ value < 0.001; *, adjusted $p$ value < 0.05; ns, not significant, adjusted $p$ value >= 0.05).

(F) Testing additional analogs for MutSβ-specific reporter activation (x-axis) and MutSα-specific reporter activation (y-axis) in K562 PEmax cells. Each small molecule was tested at a range of concentrations (0 and 20 μM) and maximal fold changes relative to DMSO controls were reported. Dots represent means of n = 3 biological replicates. Data from molecules represented in Figure 4D were also included. SAi and Si molecules that activated either of the tested reporters are indicated in red and orange, respectively. Core structures of those molecules are included (insets) with $R_1$, $R_2$, and $R_3$ representing different functional groups.

(G) Comparisons of SBi and SAi small molecules using our primary MutSβ-specific (left) and MutSα-specific (right) reporters in K562 PEmax cells. SBi and SAi compounds mainly differ from each other at two atoms in the ring scaffold, which are nitrogen atoms in SBi compounds and carbon atoms in SAi compounds. Molecules with the SBi scaffold activated the MutSβ-specific reporter more strongly than those with the SAi scaffold, while molecules with the SAi scaffold had opposite effects. Each dot represents the maximal fold change in reporter activation relative to DMSO controls for one analog, selected from a range of concentrations, as in F. Bars represent means across analog groups and error bars indicate standard deviations. Mann-Whitney U test (****, $p$ < 0.0001) was used for statistical analysis. All tested SAi and Si molecules were included in these analyses.

(H-K) Plots depict dose-dependent effects of three SAi molecules and Si-001 on MutSβ-specific reporter activation (blue lines), MutSα-specific reporter activation (red lines), and cell viability (black line, determined by Cell Titer-Glo) in K562 PEmax cells relative to the mean of DMSO controls. Dots represent fold changes in luminescence presented as means of n = 3 biological replicates and error bars indicate standard deviations. SAi and Si functional groups ($R_1$, $R_2$, and $R_3$ for SAi compounds; $R_1$ and $R_2$ for Si-001) are depicted (plot insets). Data are from the same experiments shown in F.

### FIGURE S7

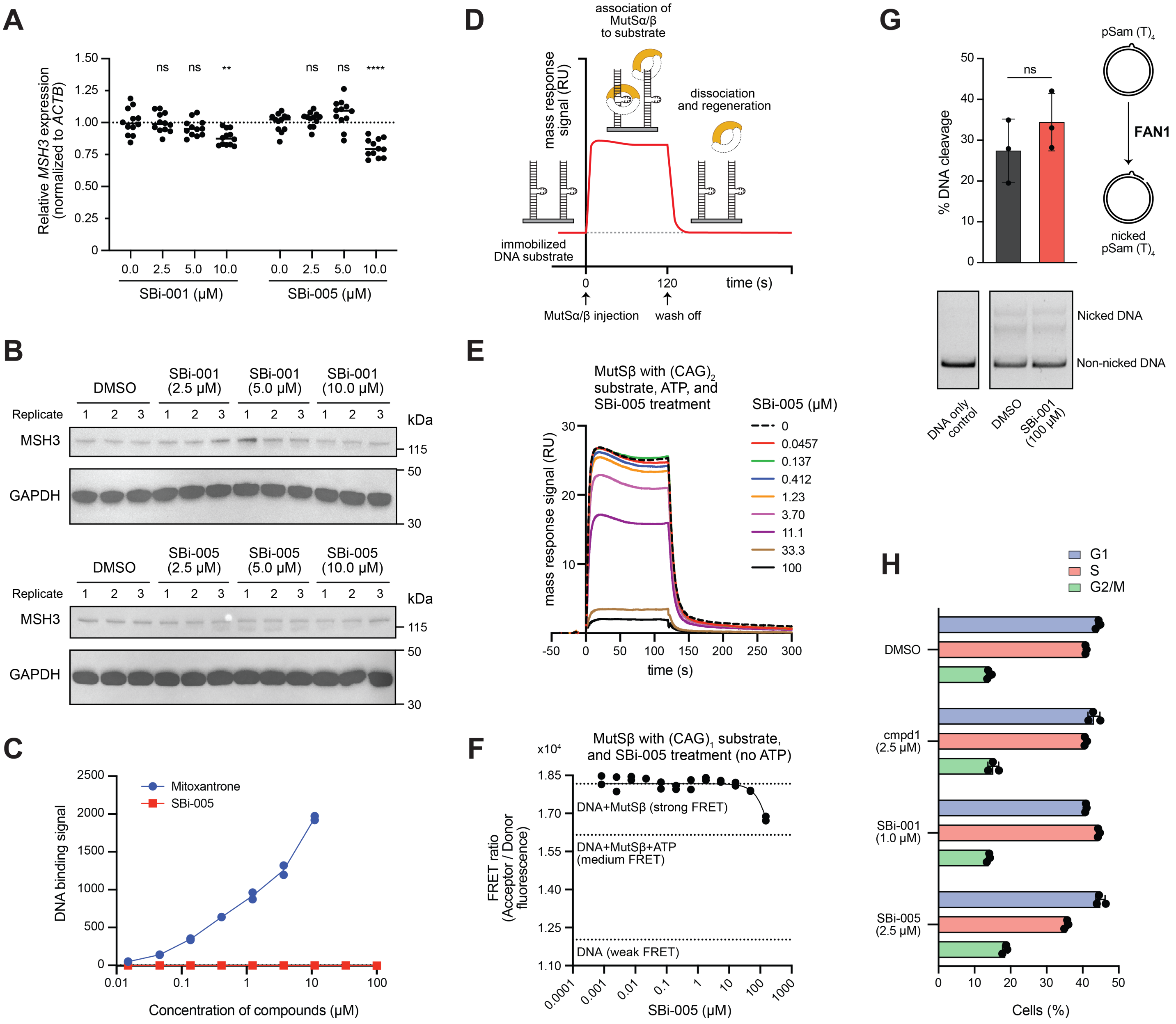

A
Relative MSH3 expression (normalized to ACTB)
ns ns ** ns ns ****
SBi-001 (μM)
SBi-005 (μM)
B
DMSO
SBi-001 (2.5 μM)
SBi-001 (5.0 μM)
SBi-001 (10.0 μM)
SBi-005 (2.5 μM)
SBi-005 (5.0 μM)
SBi-005 (10.0 μM)
Replicate
MSH3
GAPDH
kDa
C
Mitoxantrone
SBi-005
DNA binding signal
Concentration of compounds (μM)
D
association of MutSα/β to substrate
dissociation and regeneration
mass response signal (RU)
immobilized DNA substrate
time (s)
MutSα/β injection
wash off
E
MutSβ with (CAG)2 substrate, ATP, and SBi-005 treatment
SBi-005 (μM)
mass response signal (RU)
time (s)
F
MutSβ with (CAG)1 substrate, and SBi-005 treatment (no ATP)
FRET ratio (Acceptor / Donor fluorescence)
DNA+MutSβ (strong FRET)
DNA+MutSβ+ATP (medium FRET)
DNA (weak FRET)
SBi-005 (μM)
G
% DNA cleavage
ns
pSam (T)4
FAN1
nicked pSam (T)4
Nicked DNA
Non-nicked DNA
DNA only control
DMSO
SBi-001 (100 μM)
H
G1
S
G2/M
DMSO
cmpd1 (2.5 μM)
SBi-001 (1.0 μM)
SBi-005 (2.5 μM)
Cells (%)

**Figure S7. SBi molecules do not reduce expression of MSH3, intercalate in DNA, indiscriminately inhibit endonucleases, or strongly impact cell cycle progression, related to Figures 5**

(A) RT-qPCR of *MSH3* transcript from K562 PEmax cells treated with DMSO or indicated concentrations of two SBi compounds, SBi-001 or SBi-005. Dots represent relative *MSH3* expression normalized to *ACTB* control, n = 3 biological replicates with 4 technical replicates each. Brown-Forsythe ANOVA test was used for statistical analysis followed by Dunnett's T3 multiple comparison test (****, adjusted $p$ value < 0.0001; **, adjusted $p$ value < 0.01; ns, not significant, adjusted $p$ value >= 0.05).

(B) Western blot analyses of MSH3 from K562 PEmax cells treated with DMSO or indicated concentrations of SBi-001 (top) or SBi-005 (bottom). Biological replicates n = 3. Groups of images are from the same membrane, each of which was simultaneously blotted with the indicated antibodies.

(C) DNA binding of mitoxantrone (blue) and SBi-005 (red) to a 38 bp DNA containing a $(CAG)_2$ loop-out measured by SPR. Dots represent mass response signals in resonance units of n = 2 (mitoxantrone) or 1 (SBi-005) biological replicates. We note that at 33 and 100 μM mitoxantrone displayed significant binding to an empty SPR flow cell without DNA, so data from those concentrations were omitted.

(D) Schematic of SPR assay for measuring MutSα/β binding to immobilized DNA substrates with increasing concentrations of SBi-005 (Figures 5A and 5B). MutS/β was injected at 0 s and washed out at 120 s. RU indicates resonance units.

(E) Plot depicts representative data from SPR assay (as depicted in D) performed to measure MutSβ binding to a 38 bp DNA containing a $(CAG)_2$ loop-out across SBi-005 concentrations (Figure 5A). Data represented in Figure 5A are average signals from 115 to 120 s normalized to signals from no compound controls. RU indicates resonance units. Data represented in Figure 5B were similarly determined.

(F) MutSβ binding to a 70 bp DNA containing a $(CAG)_1$ loop-out at different concentrations of SBi-005 measured in the absence of ATP by FRET assay. Dots represent FRET ratios from n = 2 biological replicates and the solid black line represents a sigmoidal fit. Control signals measured without compound treatment are indicated by black dashed lines.

(G) *In vitro* cleavage of a plasmid containing a $(T)_4$ loop-out by recombinant FAN1 in the presence of DMSO or SBi-001. Dots in barplot (upper left) represent percentages of cleavage, bars indicate means of n = 3 biological replicates, and error bars indicate standard deviations. Unpaired t test with Welch's correction was used for statistical analysis (ns, not significant, $p$ value >= 0.05). Images of a representative gel (lower left, from the same gel) and a schematic of the experiment (upper right) are also included.

(H) Cell cycle analysis from a cellular model of repeat expansion (i.e., RPE-1 hTERT cells with exogenous mHTT exon 1) after 16 days of growth in the presence of DMSO or indicated compounds (vehicle and compounds replenished every 3-4 days). Dots represent percentage of cells in indicated cell cycle phase, bars represent means of n = 3 biological replicates, and error bars indicate standard deviations.
